# Host-specific plasmid evolution explains the variable spread of clinical antibiotic-resistance plasmids

**DOI:** 10.1101/2022.07.06.498992

**Authors:** F. Benz, A. R. Hall

## Abstract

Antibiotic resistance encoded on plasmids is a pressing global health problem. Predicting which plasmids spread/decline in the long term remains a huge challenge, even though some key parameters influencing plasmid stability have been identified, such as plasmid growth costs and horizontal transfer rates. Here, we show these parameters evolve in a strain-specific way among clinical plasmids/bacteria, and this occurs rapidly enough to alter the relative likelihoods of different bacterium-plasmid combinations spreading/declining. We used experiments with *Escherichia coli* and antibiotic-resistance plasmids isolated from patients, paired with a mathematical model, to show long-term plasmid stability (beyond antibiotic exposure) was better explained by evolutionary changes in plasmid-stability traits than by initial variation among bacterium-plasmid combinations. Evolutionary trajectories were specific to particular bacterium-plasmid combinations. Genome sequencing and genetic manipulation helped explain this, revealing epistatic (here, strain-dependent) effects of key genetic changes affecting horizontal plasmid transfer. Several genetic changes involved mobile elements and pathogenicity islands. Rapid strain-specific evolution can thus outweigh ancestral phenotypes as a predictor of plasmid stability. Accounting for strain-specific plasmid evolution in natural populations could improve our ability to anticipate and manage successful bacterium-plasmid combinations.

## Introduction

Plasmids are self-replicating genetic entities that play a major role in bacterial ecology and evolution. They supply their bacterial host with functional innovations, such as antibiotic- and heavy metal resistance and virulence factors (Frost *et al*., 2005; Rodríguez-Beltrán *et al*., 2021). Particularly problematic in clinical contexts are conjugative plasmids carrying antibiotic resistance genes, which can transfer horizontally (Smillie *et al*., 2010; Tamma *et al*., 2021). To manage the dissemination of resistance plasmids and associated morbidity, mortality, and healthcare costs, we need to understand the factors that drive their spread or decline in bacterial populations (Dunn *et al*., 2019; Murray *et al*., 2022). Some plasmids are associated with specific bacterial lineages, and these “successful” bacterium-plasmid combinations are a primary driver of the global dissemination of antibiotic-resistance (San Millán, 2018; Dunn *et al*., 2019). Yet it remains unclear why some combinations are more successful than others, making it difficult to predict their emergence and manage their spread. For example, the high prevalence of *E. coli* of sequence type 131 (ST131, clade C) with IncF-family plasmids encoding the extended spectrum beta lactamase (ESBL) *bla*_CTX-M_ has been ascribed to a competitive advantage during gut colonisation (McNally *et al*., 2016, 2019; Stoesser *et al*., 2016), but it is not clear why this particular bacterium-plasmid combination, and not others, has achieved this ecological success. Nevertheless, some of the key parameters affecting the ecological success of plasmids have been identified (Brockhurst and Harrison, 2021). In this paper, we ask whether the variable spread/decline of different clinical bacterium-plasmid combinations in the absence of antibiotics can be explained by variation of these parameters, both among different combinations and over evolutionary time.

Plasmid persistence is generally expected to depend on the balance of growth costs (how plasmids affect bacterial population growth) and transfer rates (typically by conjugation) and, to a lesser extent given many natural plasmids carry addiction systems, segregational loss (Stewart and Levin, 1977; Levin *et al*., 1979; Finbarr, 1995; Ponciano *et al*., 2007; San Millán *et al*., 2014). Past work showed growth costs (Alonso-del Valle *et al*., 2021; Dunn *et al*., 2021), transfer rates (Lopatkin *et al*., 2017; Sheppard *et al*., 2020) and segregational loss (De Gelder *et al*., 2007) can vary depending on the bacterium-plasmid combination. This variability may therefore be central to predicting the long-term stability of plasmids (over hundreds of generations and in the absence of antibiotics). However, bacteria and their plasmids can evolve rapidly, for example to reduce growth costs or alter transfer rates, thereby promoting plasmid stability (Turner *et al*., 1998; Dahlberg and Chao, 2003; De Gelder *et al*., 2008; Harrison *et al*., 2015, 2016; Loftie-Eaton *et al*., 2016, 2017; Porse *et al*., 2016; Dimitriu *et al*., 2021). Such evolutionary changes may reduce the predictive power of plasmid-stability traits measured at a given point in time. In particular, if different bacterium-plasmid combinations vary in their propensity for evolutionary changes in plasmid-stability traits, then initial variation of those traits may be a poor predictor of which combinations are most stable in the long term. However, whether and how evolution of plasmid-stability traits varies among bacterium-plasmid combinations remains poorly understood. Despite this, epistasis has been documented for chromosomal mutations affecting plasmids (San Millán *et al*., 2015; Loftie-Eaton *et al*., 2016), indicating their evolutionary trajectories may indeed be specific to individual combinations. In summary, we know plasmids and bacteria evolve rapidly, but is this fast, strong and variable enough to ‘reshuffle the deck’ and change which bacterium-plasmid combinations are most stable in the long term?

Here, we test whether the spread/decline of resistance plasmids in experimental populations of *E. coli* strains over 15 days (∼150 generations) in the absence of antibiotics is explained by (i) initial variation of plasmid-stability traits among bacterium-plasmid combinations and/or (ii) variable evolution of plasmid-stability traits during the experiment. We generated multiple bacterium-plasmid (hereafter, strain-plasmid) combinations from clinical *E. coli* strains and their natively associated ESBL-plasmids. This approach overcomes some limitations of previous research into the role of rapid evolution in the spread of plasmids, which has typically focused on one strain at a time (Stalder *et al*., 2017; Hall *et al*., 2021), indirect genomic evidence from bacteria evolving in clinics (Knudsen *et al*., 2018; Lu *et al*., 2020; León-Sampedro *et al*., 2021), or model strains/plasmids (San Millán *et al*., 2014; Loftie-Eaton *et al*., 2017). Using clinical strains/plasmids is particularly important here, first because they are more relevant for the resistance crisis, and second because genetic factors, such as other plasmids and virulence determinants that are lacking from laboratory strains, may play key roles in the spread of the focal resistance plasmids we aim to understand (Benz *et al*., 2020; Haudiquet *et al*., 2021). We tracked the frequencies of ESBL-plasmid carrying clones during serial passage in the absence of antibiotics and quantified plasmid-stability traits of ancestral and evolved clones. Implementing these as parameters in a mathematical model, we simulated plasmid dynamics for each strain-plasmid pair. With analyses of whole-genome sequences we identified parallel, yet strain-plasmid-pair-dependent, evolution. Together, this shows how rapid plasmid evolution can override initial variation among strain-plasmid pairs and reveals its dependence on the specific strain-plasmid combination, as exemplified by epistatic plasmid mutations increasing plasmid transfer rates.

## Results

### Stability of resistance plasmids varies among strain-plasmid combinations

We generated six unique strain-plasmid pairs, each derived from one of three *E. coli* strains isolated from hospital patients and one of two ESBL-encoding clinical resistance plasmids natively associated with the same strains (**Fig. 1**). To monitor the spread/decline of each ESBL-plasmid in populations of each strain, we made replicate microcosm cultures, each containing a mixture of a plasmid-carrying strain and the corresponding plasmid-free strain. We then serially passaged each population for 15 days in the absence of antibiotics. We used the frequency of resistance phenotypes as a proxy for the frequency of ESBL-plasmid carrying clones over time, and verified this reflected plasmid carriage by PCR screening (see Supplementary Methods; **Tables S1-S2**). In all populations, ESBL-plasmids persisted over 15 days of antibiotic-free serial passage (**Fig. 2**). However, the temporal plasmid dynamics varied among strains and plasmids (two-way ANOVA, effect of plasmid: *F*_1,17_ = 14.758, *p* < 0.01; effect of strain: *F*_2,17_ = 7.834, *p* < 0.01; the response variable here is the area under the curve, AUC, of plasmid frequency over time, a measure of time-averaged plasmid success). The differences among plasmids also depended on which strain they were carried by (plasmid × strain interaction: *F*_2,17_ = 6.672, *p* < 0.01). For example, both plasmids spread rapidly in populations of strain 15, but showed different trajectories in populations of strain 19 (**Fig. 2**). This difference in long-term stability between the two plasmids is surprising given their high sequence similarity (**Fig. S1**). One possible explanation would be an average difference in their initial frequencies in microcosms containing the different strains, but we excluded this in a further experiment (**Fig. S2**). There was also marked variation among replicate populations in some combinations, such as those with strain 1 carrying either plasmid **(Fig. 2**).

**Figure 1:**
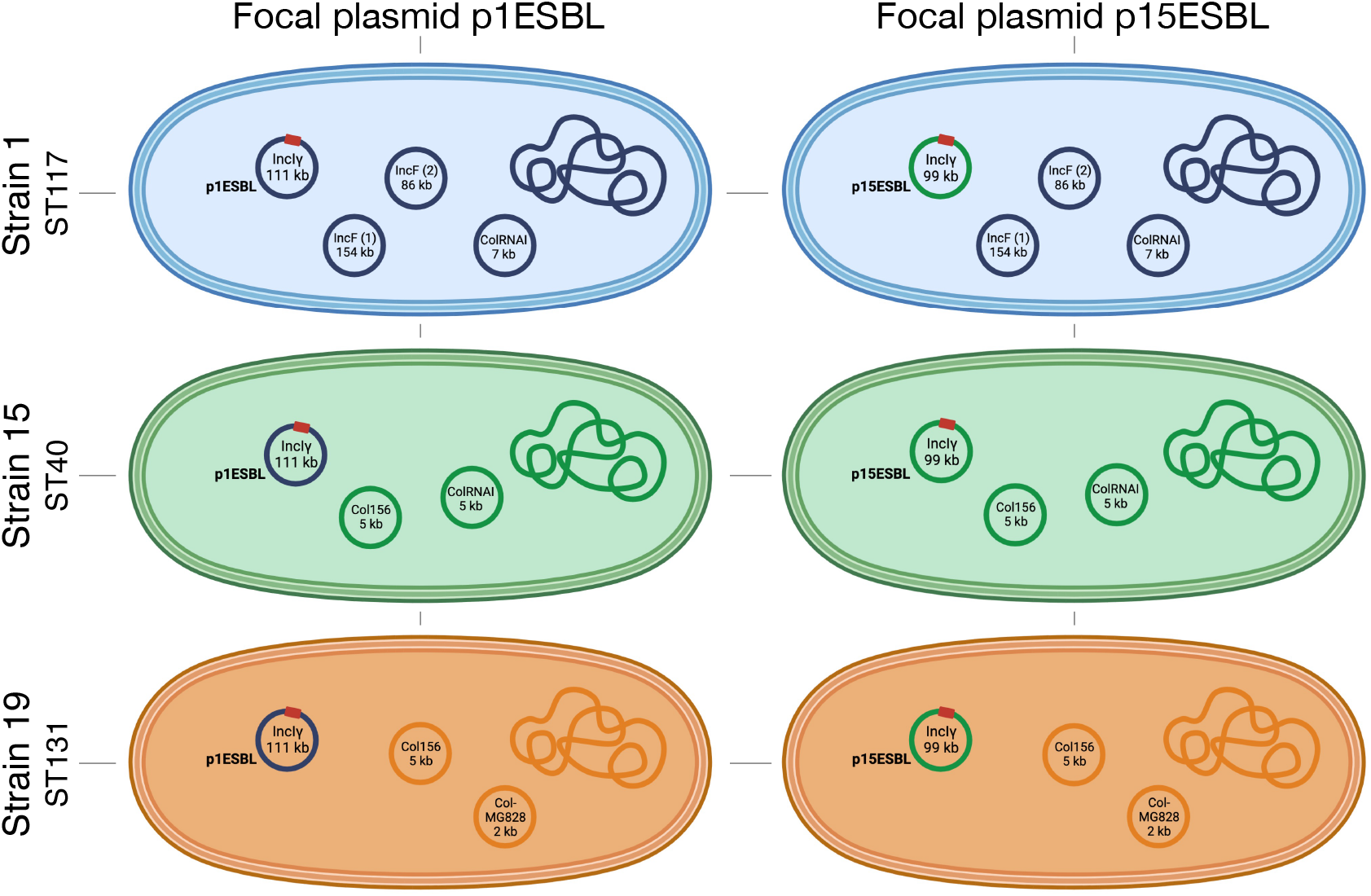
Six unique strain-plasmid pairs. Clinical *E. coli* strains (1, 15 and 19; rows) each carrying one of two focal plasmids (p1ESBL, an ESBL-plasmid originally from strain 1, or p15ESBL, an ESBL-plasmid originally from strain 15; columns). The red square represents an ESBL-encoding gene on the focal plasmid in each strain; ST gives the sequence type of each strain. In cases where plasmids carry multiple replicons, only one is shown. See **Table S3** for detailed description of plasmids and their antibiotic-resistance genes.

**Figure 2:**
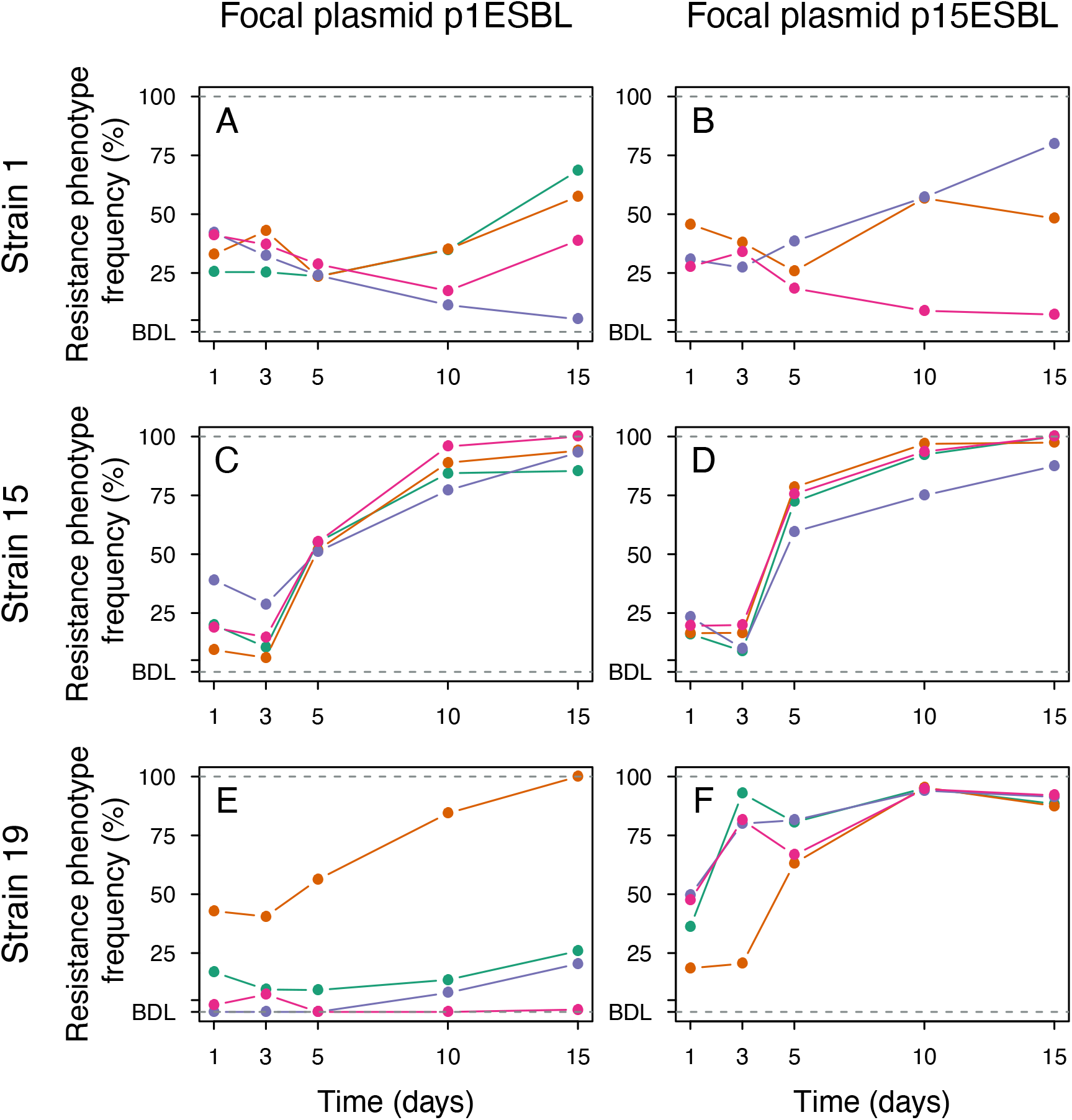
Strain-plasmid-pair-dependent plasmid dynamics in the absence of antibiotics. Panels **A-F** show the different strain-plasmid pairs. In each panel, colors indicate the four replicate populations (three in B) and are maintained in the other figures. The y-axis gives the fraction of colony-forming units with an ESBL phenotype (approximating the frequency of plasmid-carrying cells, which we confirmed by PCR; see Supplementary Methods) in each replicate population sampled at days 1, 3, 5, 10 and 15. Points below the detection limit (BDL) indicate cases where no ESBL-positive cells were recovered during replica-plating.

### Variable plasmid dynamics are poorly explained by initial variation of plasmid-stability traits

We estimated plasmid-stability traits for each ancestral strain-plasmid pair (**Fig. 3A-B**). Growth costs (population growth rate of a plasmid-carrying strain relative to its plasmid-free equivalent) varied depending on the bacterial strain, but not which of the two plasmids it carried (two-way ANOVA, effect of strain: *F*_2,26_ = 10.031, *p* < 0.001; effect of plasmid *F*_1,26_ = 0.114, *p* = 0.739; **Fig. 3A**), with particularly large costs for both plasmids with strain 19. Transfer rates varied depending on the plasmid, the host strain, and their interaction (two-way ANOVA, plasmid effect: *F*_1,12_ = 76.620, *p* < 0.001; strain effect *F*_2,12_ = 6.627, *p* < 0.05; plasmid × strain interaction *F*_2,12_ = 19.360, *p* < 0.001; **Fig. 3B**). For example, transfer rate of p1ESBL in strain 19 was very low compared to p15ESBL, whereas both plasmids transferred similarly well in strain 15. We did not detect significant plasmid loss for any strain-plasmid pair over 24 hours (**Fig. S3**), which we attribute to the toxin-antitoxin (TA) systems present on both plasmids (see Methods). Qualitatively, this variation in ancestral plasmid-stability traits (**Fig. 3A-B**) is consistent with some of the observed plasmid dynamics over 15 days (**Fig. 2**; **Fig. 3C**): both plasmids spread rapidly with strain 15 and these combinations had relatively low costs and high transfer rates. However, other long-term trends were not consistent with the ancestral variation of plasmid-stability traits, such as the rapid spread of p15ESBL with strain 19 (**Fig. 2**; **Fig. 3C**) despite this combination having the second largest fitness cost (**Fig. 3A**).

**Figure 3:**
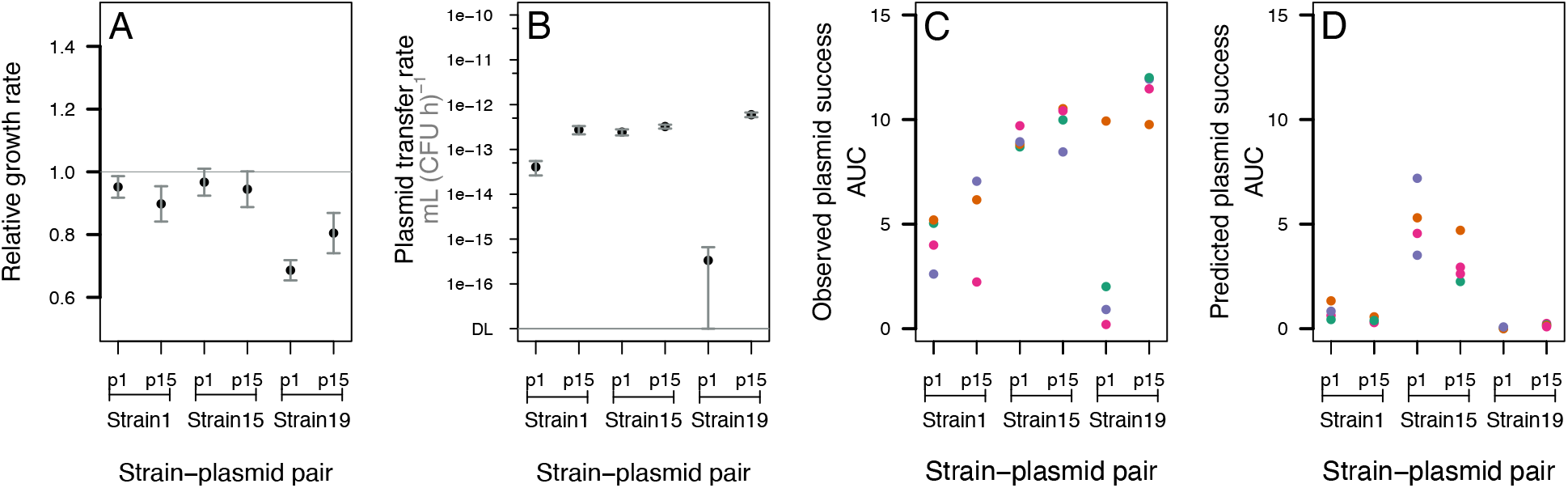
Ancestral plasmid-stability traits explain only some of the observed plasmid dynamics. **A)** Plasmid growth costs, expressed as population growth rate of plasmid-carrying relative to equivalent plasmid-free strains in the absence of antibiotics. **B)** Plasmid transfer rates of each ancestral strain-plasmid pair, measured in mating assays with each corresponding plasmid-free equivalent as the recipient strain (see Methods). **C)** Plasmid success, estimated as the area under the curve (AUC) of plasmid frequency over time, for each strain-plasmid pair during the 15-day experiment shown in **Fig. 2. D)** Plasmid success predicted by a model simulating plasmid frequencies over time, based on parameter values from panels A and B (hereafter, the ancestral (Anc) model), with C=1.15e12 (see Methods).

We next tested the predictive power of the ancestral plasmid-stability traits quantitatively, using a variation of the well-established population dynamics model by Simonsen et al. (1990), modified by Huisman et al. (2022) to account for variable growth costs and transfer rates (see Supplementary Model Information sections I-V; **Figs. S4-S8**). Consistent with observed dynamics (**Fig. 2**; **Fig. 3C**), the model predicted both plasmids to spread effectively in populations of strain 15 (**Fig. 3D**). With strain 19, the model matched the observed decline of p1ESBL, but not the high frequency of p15ESBL (**Fig. 3C-D**). At the level of individual replicates, which are accounted for in the model by including the initial subpopulation densities specific to each replicate, the model failed to predict observed variation among replicates of strain 1 with both plasmids and strain 19 with p1ESBL. An alternative version of the model that resulted in more realistic dynamics in terms of total population growth (but not plasmid dynamics) produced a very similar outcome (see Supplementary Model Information section III). In summary, ancestral plasmid-stability traits varied among strain-plasmid pairs, but this information explained only some of the observed plasmid dynamics.

### Phenotypic evolution varies among strain-plasmid pairs and explains variable plasmid stability

To test for evolutionary changes in plasmid-stability traits, we first measured population growth rates of evolved (day 15) and ancestral (day 0) ESBL-plasmid-carrying (p^+^) and plasmid-free (p^-^) clones (**Fig. 4A**). For all strain-plasmid pairs, growth rates of evolved p^+^ clones were higher than corresponding ancestral p^+^ clones. In some populations, evolved p^+^ clones also had higher growth rates than evolved p^-^ clones from the same population, but this varied among strain-plasmid combinations (ANOVA with growth rate of evolved p^+^ relative to evolved p^-^ as the response variable: plasmid × strain interaction: *F*_2,16_ = 6.453, *p* < 0.01). For example, evolved p15ESBL^+^ clones were consistently slower growing than evolved p^-^ clones from the same populations with strains 1 and 15, but not with strain 19. This indicates growth costs of plasmids, both relative to ancestral clones and relative to co-existing plasmid-free clones, were reduced to different degrees depending on strain-plasmid combination. For p^-^ control populations (pFREE in **Fig. 4A**), evolved clones from all strain-plasmid pairs grew faster than equivalent ancestral p^-^ clones, showing there was also plasmid-independent adaptation in these conditions.

**Figure 4:**
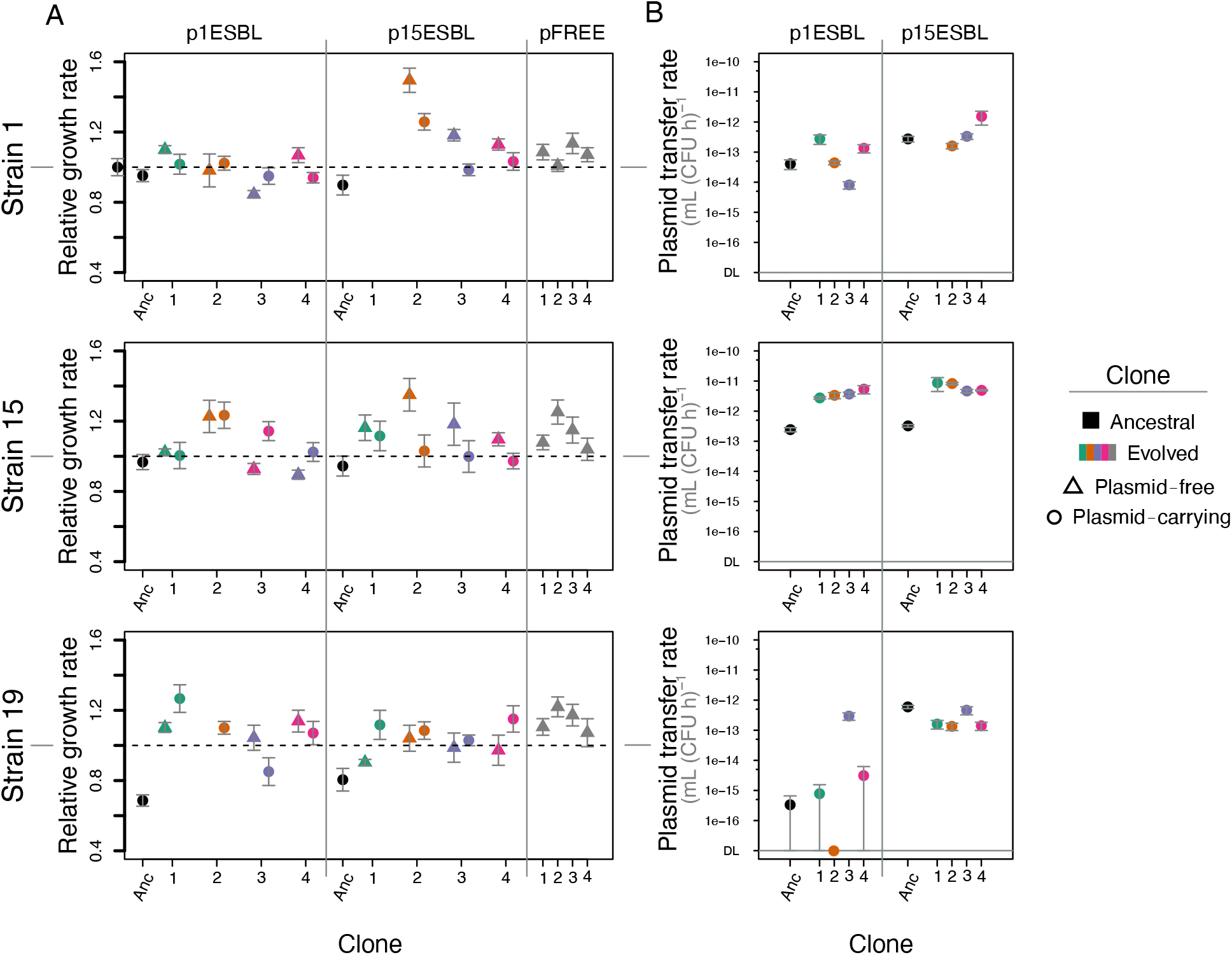
Strain-plasmid-combination-dependent evolutionary changes. **A)** Population growth rates for ancestral plasmid-carrying, evolved plasmid-carrying and evolved plasmid-free clones (see legend/x-axis) for each strain-plasmid combination, and for evolved clones from plasmid-free control cultures (pFREE at the right of each panel). Growth rate in each case is shown relative to the corresponding plasmid-free ancestral strain. **B)** Plasmid transfer rates for ancestral and evolved plasmid-carrying clones (see legend/x-axis) in each strain-plasmid combination. Transfer rates were measured in mating assays with the corresponding ancestral plasmid-free strain as the recipient. For one replicate population with strain 19-p1ESBL (orange), we found no ampicillin-susceptible colonies at the end of the experiment, so only the plasmid-carrying evolved clone is shown.

To test for evolutionary changes in plasmid transfer rates, we performed mating assays between evolved/ancestral p^+^ clones and their ancestral p^-^ equivalents (**Fig. 4B**). This revealed strong strain-dependence: both plasmids increased their transfer rates in all replicate populations with strain 15 (∼10-20 fold for p1ESBL and 20-35 fold for p15ESBL), but much less consistently and to a smaller extent with the other two strains (aligned ranks transformation ANOVA with transfer rate of evolved clones relative to ancestral as response variable, strain effect: *F*_2,16_ = 6.2942, *p* < 0.01). Changes in transfer rate also depended on the plasmid itself and the strain-plasmid combination (plasmid effect: *F*_1,16_ = 8.4926, *p* = 0.01; plasmid × strain interaction: *F*_2,16_ = 3.7065, *p* < 0.05). We found no evidence variation of plasmid success over 15 days was linked to variable segregational plasmid loss among different evolved clones within each strain-plasmid combination or on average among the different combinations (measured over 24 h; **Fig. S9**). Thus, both growth costs and transfer rates changed during our experiment, in ways dependent on the strain-plasmid combination.

To test whether the phenotypic changes specific to strain-plasmid-combinations observed above explained the variable long-term stability of plasmids (**Fig. 2**), we extended our population dynamics model to include evolved plasmid-stability traits (Evo Model). We simulated all possible combinations of ancestral/evolved bacteria (B_a_/B_e_) and plasmids (P_a_/P_e_), resulting in strains B_a_, B_a_P_a_, B_a_P_e_, B_e_, B_e_P_a_, B_e_P_e_ (see Supplementary Model Information section II; **Fig. S5**), and assumed mutant (evolved) strains/plasmids to be present at initially very low frequencies. This model revealed evolutionary changes in plasmid transfer rate and growth costs to be a better predictor of long-term plasmid stability than the ancestral phenotypes: the updated model explained more of the experimentally observed variation of plasmid stability (simulated vs observed plasmid success: *R*^2^ = 0.31, *p* < 0.05 for Evo Model 1; **Fig. 5A-B**). The Evo and Anc Models even predicted different rank orders in terms of which strain-plasmid combinations would be most successful. For example, Evo Model 1 predicted p15ESBL to spread more successfully with strain 19 than the Anc Model did (**Fig. 5A-B**), and this was much closer to what we observed in our 15-day experiment. Together, these results suggest rapid evolution of plasmid-stability traits can change which plasmid-strain combinations are most successful, thereby “reshuffling the deck” and manifesting the role of evolution in identifying successful strain-plasmid pairs.

**Figure 5:**
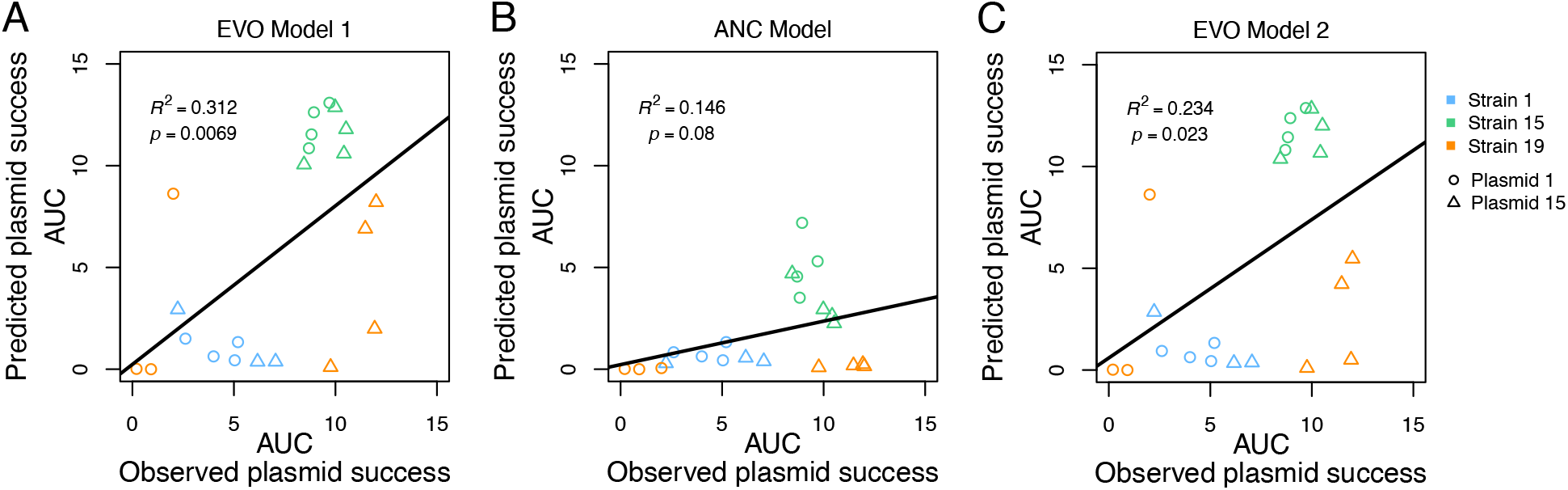
Evolutionary changes in plasmid-stability traits explain long-term plasmid stability better than ancestral values do. Plasmid success over time (AUC) observed in our 15-day serial passage experiment, x-axis in each panel, is plotted against values predicted by three different versions of the population dynamic model, panel A-C, for each replicate of each strain-plasmid combination (see legend). **A)** Evolved model 1, accounting for plasmid-stability traits of evolved clones. **B)** Ancestral model accounting for ancestral plasmid-stability traits only (as in **Fig. 3D**). **C)** Evolved model 2, accounting for evolved plasmid-stability traits, but assuming plasmid growth costs in evolved strains B_a_P_e_, B_e_P_a_ determined by the bacteria rather than the plasmid. See Supplementary Model Information for model details. Note replicate population 3/orange of strain 19-plasmid 1 is excluded in comparisons of ANC and EVO models because no susceptible evolved clones were isolated from that population (see Methods).

The superior fit of the Evo Model 1 compared to the Anc Model might be expected simply because it is more complex (wider range of subtypes). However, when we used the same Anc Model but with parameters from evolved strains (that is, without changing the number of subtypes), we obtained similar explanatory power as in the more complex Evo Model 1 (*R*^2^ = 0.31, *p* < 0.05; Supplementary Model Information section IV; **Fig. S7**). Thus, the improved fit of the Evo Model was not simply due to its complexity, but rather due to the properties of evolved strains. Also, the Evo Model included some subtypes for which we did not have experimentally determined growth rates (B_a_P_e_, B_e_P_a_; Supplementary Model Information section II). Above we assumed growth costs of plasmids in these cases to be determined by the plasmid itself. An alternative model where we instead assumed these growth costs were determined by the bacterial host (Evo Model 2) supported the same conclusions (simulated vs observed plasmid success: *R*^2^ = 0.23, *p* < 0.05; **Fig. 5C**). Finally, we investigated the sensitivity of our conclusions to a key assumption of our model and those it is based on (Simonsen *et al*., 1990): that conjugative transfer does not take place in stationary phase. In some model versions, relaxing this assumption increased average predicted plasmid succces and explained even more of the observed variation in plasmid dynamics (Supplementary Model Information section V; **Fig. S8**). This last result was only the case when we assumed both unrealistic total population growth and continuous plasmid transfer by mass action, and should therefore be interpreted with caution.

### Parallel genetic changes in ESBL-plasmids and chromosomes are specific to the strain-plasmid combination

We whole-genome sequenced ancestral strain-plasmid combinations (short- and long read-sequencing (Benz *et al*., 2020; Huisman *et al*., 2021)) and single endpoint clones (p^+^/p^-^, Illumina only) from each replicate population. First, we found each evolved clone maintained all their native non-ESBL plasmids. Thus, we could exclude changes in carriage of non-focal plasmids as a driver of observed variation in ESBL-plasmid dynamics (**Fig. 2**). Further, we found plasmid ColRNAI from strain 1 had co-transferred to strain 19 when generating strain19-p1ESBL, where it was maintained in two out of the 4 sequenced ESBL-plasmid-carrying evolved clones (green and orange). Sequence data also revealed strain-specific parallel evolution (**Fig. 6**). In combination with strain 15, both ESBL-plasmids showed parallel genetic changes in every independently evolved replicate clone (same plasmid region, different nucleotide changes; **Fig. 6B**). These genetic changes on plasmids were specific to strain 15 (not found in other evolved clones; **Fig. 6**), and coincided with increased plasmid-transfer rates in these clones (**Fig. 4**).

**Figure 6:**
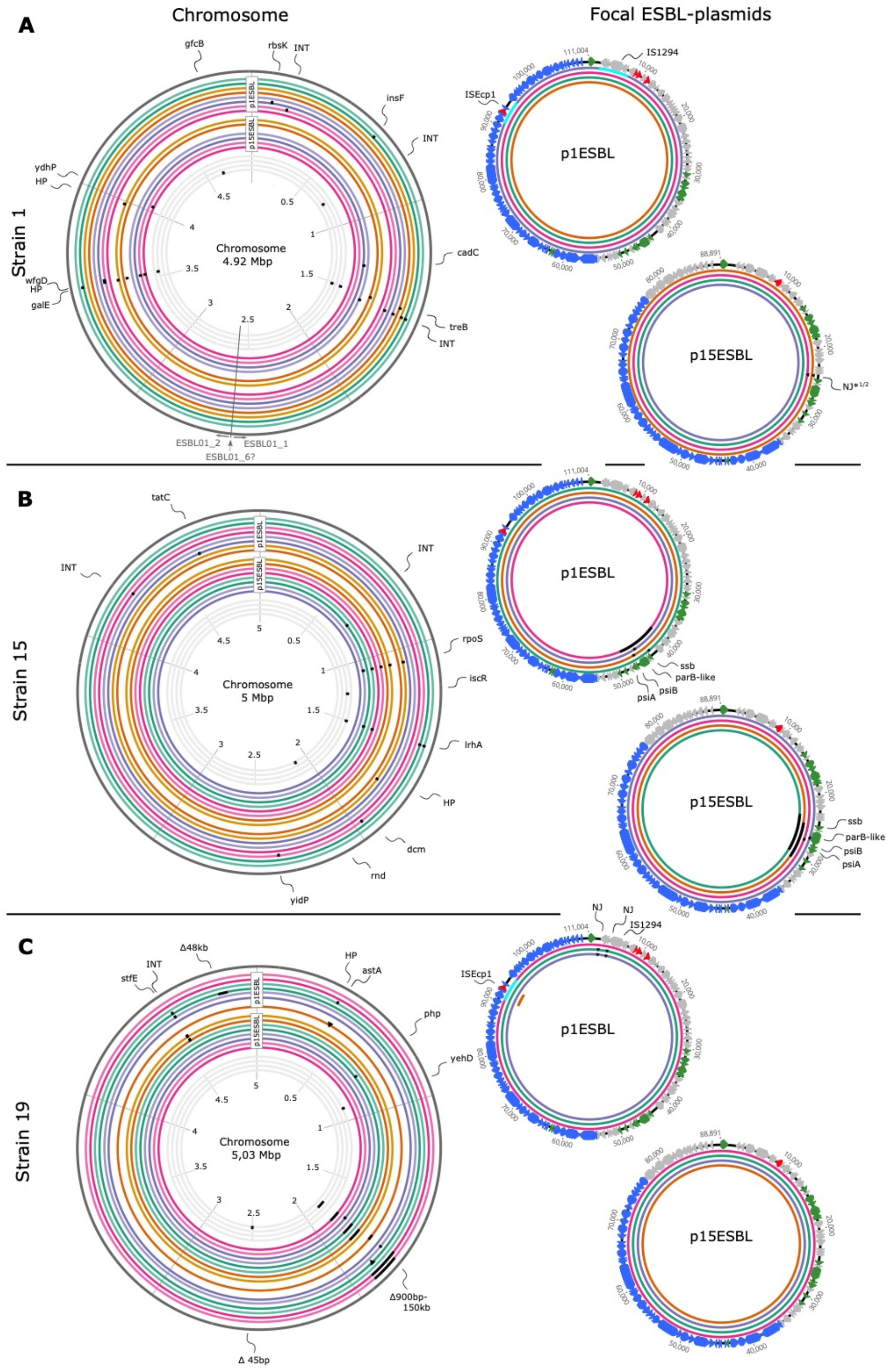
Genetic changes in chromosomes and ESBL-plasmids after 15 days of serial passage without antibiotics. Chromosomes (left) are shown for each strain (**A-C**) evolved with either of the two focal ESBL-plasmids (shown at right). In chromosome plots, the eight outermost circles are clones isolated from populations evolved with p1ESBL, the eight middle circles are clones isolated from populations with p15ESBL, and the four innermost circles are clones from plasmid-free control populations. Colours indicate replicate populations; for each population two evolved clones are shown (darker shade = plasmid-carrying; lighter shade = plasmid-free). Black squares show non-synonymous genetic changes and arrows indicate target genes/regions. In the chromosome, only deletions of 20 kb and larger are to scale, triangles indicate new junctions (NJ) and turquoise squares inversions of that region. In the plasmid plots, colored rings identify replicate populations (as in chromosome plots). Plasmid genes are colored according to predicted function: red = antibiotic resistance, blue = transfer genes, green = stability genes, grey = other. Non-focal plasmids did not show genetic changes and are not depicted. **Table S9** gives a detailed list of genetic changes.

In several cases, we detected genetic changes involving interactions between loci on ESBL-plasmids and other mobile genetic elements, including mobile elements encoded on the ESBL-plasmids themselves (**Tables S4-S5**). For example, in the ancestral p1ESBL, the shufflon segment B is disrupted by a 2880 bp transposition unit, encoding the insertion sequences ISEcp1 and the *bla*_CTX-M-1_ gene (**Fig. S10**). In one sequenced clone each of strain 19 and strain 1, the shufflon segments of p1ESBL rearranged such that the entire transposition unit was inverted (**Fig. 6A/C**, turquoise squares). In these two populations, p1ESBL was at very low frequency at day 15 (**Fig. 2)**, and showed decreased transfer rates in combination with strain 1 (**Fig. 4**). In a strain 19 clone from another replicate population, the same mobile element jumped in a cut-and-paste manner to the chromosome (**Fig. 6C** (*astA*); **Fig. S10**) while the same clone lost p1ESBL, explaining the non-transferable resistance phenotype of this evolved clone (**Fig. 4**). The other plasmid, p15ESBL, also showed evidence of interactions among genetic elements: p15ESBL from two strain 1 clones from different replicate populations had a new genetic junction (NJ, triangles in **Fig. 6**; here NJ*^1/2^ in **Fig. 6A**) with a putative group II intron. This intron is also present in the ancestral IncF-plasmid and p1ESBL of strain 1 (ESBL01_04913, **Table S6**). Furthermore, this region in p1ESBL is identical to the position of NJ*^1/2^ in p15ESBL, and is adjacent to the mutated locus on ESBL-plasmids with strain 15 described above (**Fig. 6B**). Thus, several genetic changes and interactions with other mobile elements occurred in this region of our plasmids. Finally, we detected evidence of moderately increased copy number for evolved vs ancestral plasmids, with one strain 19-plasmid 1 replicate showing a particularly large increase (**Fig. S11**).

Genetic changes in bacterial chromosomes also demonstrated a high level of parallelism and, as above for plasmid mutations, often involved accessory genomic features specific to clinical or pathogenic bacteria, such as pathogenicity islands (PAIs). Each sequenced strain 19-p15ESBL clone (from independent populations) had a 900 bp -150 kb deletion of or within a putative chromosomal PAI (**Fig. 6C**; **Table S7**). These deletions were specific to the p^+^- clones, which consistently had a growth advantage over the corresponding p^-^- endpoint clones (**Fig. 4**). Also in strain 1, ∼20 kb of a putative chromosomal PAI (**Fig. S12**; **Table S8**) was deleted in 8 out of the 19 sequenced evolved clones but never in clones carrying p15ESBL. We found parallel mutational changes in *rpoS* (central regulator of the general stress response) specific to p^-^- endpoint clones only in populations of strain 15 where p15ESBL (and in one case p1ESBL) was present. These mutations were either frameshift mutations or SNPs introducing a preliminary stop codon and all these evolved p^-^-clones could outgrow the evolved p15ESBL-carrying clones from the same populations (**Fig. 4**). In summary, chromosomal evolution depended on both the host strain and which ESBL-plasmid was present, and using clinical strains here allowed us to detect adaptive changes in genetic elements such as PAIs that are often not represented in laboratory or model strains.

### Epistatic changes in plasmid leading region explain strain-specific increases in transfer rates

Having found above that evolved p1ESBL and p15ESBL in strain 15 increased their transfer rates, and acquired parallel genetic changes, we asked why these mutations were not observed in the other strains. These genetic changes were in the plasmid leading region, the first part of a conjugated plasmid to enter the recipient cell and expressed immediately thereafter in single-stranded form via single-strand initiation promoters (*ssi*, (Bates *et al*., 1999; Virolle *et al*., 2020)). Observed changes were likely loss-of-function mutations: they included SNPs introducing a preliminary stop codon, insertions leading to a frame shift, deletions within one operon or its promoter *ssi3*, or the entire operon (**Fig. 7A**). To test whether genetic changes in this region confer altered plasmid transfer rates, we generated two different knock-out mutants disrupting the *ssi3* operon; mutA (Δ*parB*-like gene) and mutB (Δ*parB*-like gene, Δ*psiB*, Δ*psiA*) in both ancestral plasmids (p1ESBL and p15ESBL; **Fig. 7B**). In combination with strain 15, mutA and mutB both increased the transfer rates of both ESBL-plasmids (**Fig. 7C**), suggesting this region does indeed affect transfer rate. However, this was not the case with strain 1, demonstrating the effect of these mutations is strain-specific (**Fig. 7C**; Welch Two Sample t-test for strain 15, *p* < 0.05 in all cases before and after Holm’s correction for multiple testing; Wilcoxon Rank Sum Test for strain 1, *p* > 0.05 in all cases before and after Holm’s correction for multiple testing). Introducing mutA or mutB did not affect plasmid growth cost in strain 1 or strain 15 (**Fig. S13**). Strain 19 is not included here because we could not re-introduce mutant plasmids into the ancestral plasmid-free strain 19 (see Methods). In summary, this plasmid locus, showing parallel strain-specific evolution in our 15-day experiment and coinciding with altered transfer rates, affected plasmid transfer in a strain-specific way. This is consistent with a strain-specific benefit of these plasmid mutations in terms of boosting transfer, and therefore plasmid abundance, in the absence of antibiotics. Moreover, that the same region was mutated in both plasmids suggests mutations in the *ssi3* operon may be a general mechanism of adaptation in clinically relevant plasmids such as these.

**Figure 7:**
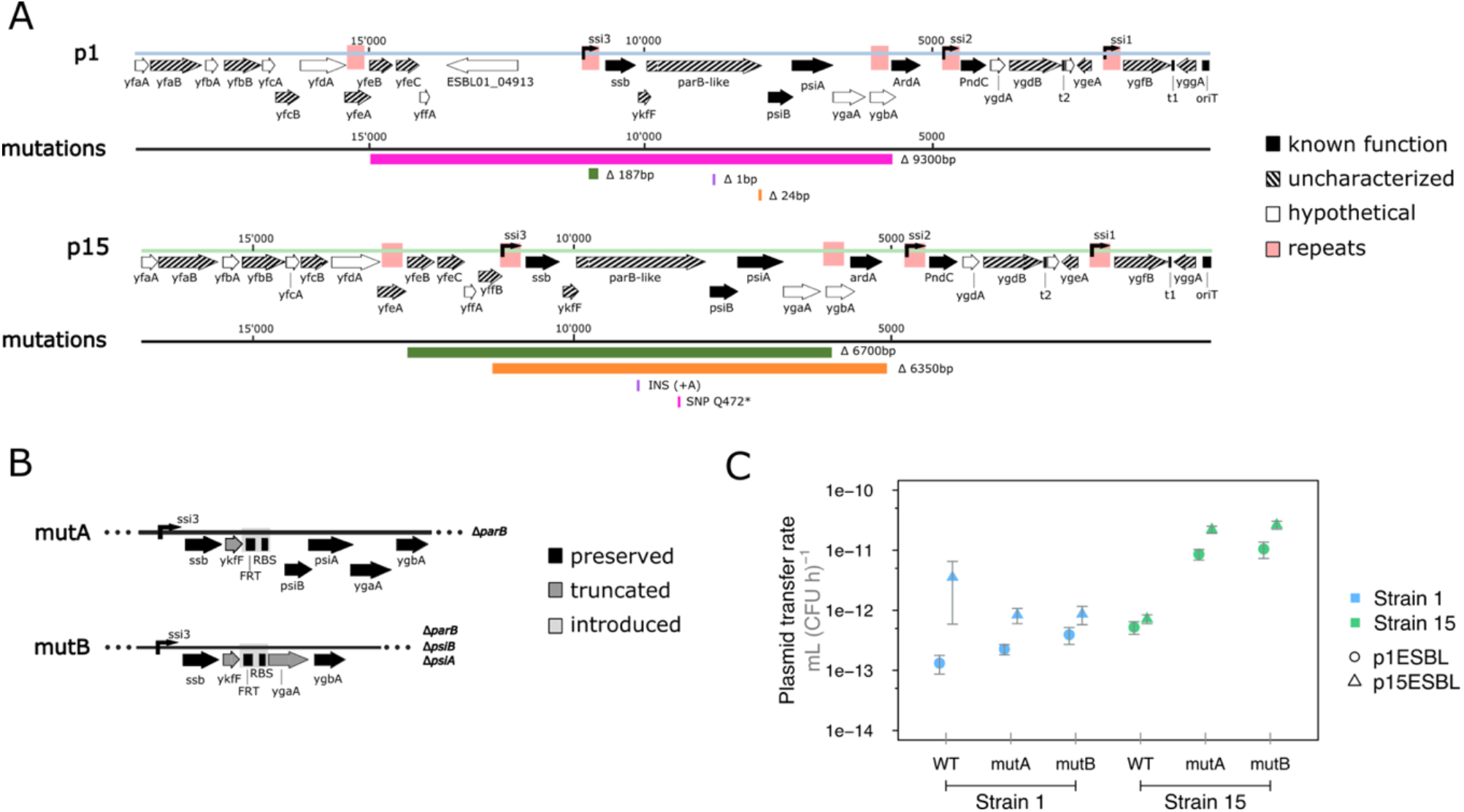
Genetic changes in the plasmid leading region increase transfer rate in a host-specific way. **A)** Plasmid leading region (oriT to ∼12 kb downstream) of p1ESBL and p15ESBL and mutations accumulated during the evolution experiment. This region consists of three operons, transcribed by ssi3-1, with the following genes: ssb; ssDNA binding protein, parB-like gene; unknown function, psiB-psiA; ssDNA evoked SOS-response inhibition, ardA; anti restriction protein, PndC; post-segregation killing protein, and several open reading frames (ORFs) of unknown function. Further information about genes/functions is in **Table S6**. Mutation colours correspond to the replicate clone they were found in. **B)** Schematic of the ssi3 operon for the knock-out mutants mutA and mutB. RBS stands for ribosomal binding site and FRT for flippase recognition target, which were introduced during mutant construction. Both mutants have a truncated ykfF and likely exhibit polar effects on expression of the downstream genes in the ssi3 operon (psiB, psiA, ygaA, ygbA). **C)** Both mutant plasmids increased transfer rates with strain 15 only.

## Discussion

To manage the global spread of antibiotic resistance, we need to understand which bacterium-plasmid combinations are most successful. Plasmid persistence in the absence of antibiotics has been shown to depend on plasmid stability traits (growth cost, horizontal transfer and segregational loss), which vary among different bacterium-plasmid combinations (Lopatkin *et al*., 2017; Benz *et al*., 2020; Alonso-del Valle *et al*., 2021; Hall *et al*., 2021). Our results show evolutionary changes of these parameters can outweigh their initial variation among strain-plasmid combinations in explaining plasmid persistence, rapidly altering both absolute and relative stability across different combinations. For example, the strain-plasmid combination with the second-largest initial growth cost in our experiment was one of the most successful in the long term (strain 19-p15ESBL). Using a quantitative model to predict the relative stabilities of different strain-plasmid combinations, we gained more explanatory power (both quantitatively and qualitatively, in terms of the rank order of our predictions) by accounting for evolutionary changes occurring over 15 days. Strain-specific evolutionary changes included mutations affecting plasmid transfer rate, which we observed consistently with one of our strains but not with others. Genetic manipulation experiments demonstrated mutations in this region have epistatic (here, strain-specific) effects on transfer rate. Thus, rapid evolution can “reshuffle the deck” and here played a greater role than initial variation among strain-plasmid combinations in directing the trajectories of clinically important plasmids in the absence of antibiotics.

Our finding of a strong role for rapid evolution here is particularly relevant for predicting the spread of resistance because we used clinical bacterial strains and plasmids (San Millán, 2018). Clinical strains naturally carry other plasmids, accessory regions such as PAIs, phages (**Table S10**) and other mobile elements (Hülter *et al*., 2017; Rodríguez-Beltrán *et al*., 2021). These features may significantly alter the evolutionary trajectory of plasmids and their hosts compared to lab strains or model plasmids. In support of this approach revealing new insights into plasmid evolution, we detected several genetic changes at several such loci, often involving interactions within and among mobile elements, such as intron and transposon movements (**Fig. 6**) and the loss of PAIs (**Fig. 6; Fig. S12**). Crucially, key mechanisms of adaptation in our experiment (recombination/mutation in the leading region; loss of PAI islands) have also been observed in nature (Schmidt and Hensel, 2004; Venturini *et al*., 2013). Furthermore, some of the most important genetic changes in our experiments occurred at the same loci on both plasmids. This suggests the rapid evolutionary changes in our experiments may apply to other plasmids and in other settings, both in the context of antibiotic resistance and the wider role of plasmids in bacterial evolution.

Our results provide new insights into genetic mechanisms stabilising plasmids during evolution. Parallel evolution of both ESBL-plasmids revealed the leading region as an important evolutionary target. This region is conserved (Virolle *et al*., 2020) (**Table S6**), but variants differ by mobile elements such as transposable or IS elements (Takahashi *et al*., 2011). Within the leading region, we consistently found changes affecting the gene encoding a ParB-like hypothetical protein (**Table S6**) in evolved clones of strain 15 with both plasmids. These included frameshifts, SNPs and deletion of the entire *ssi3* operon (**Fig. 7A**). These deletions often involved recombination of repeat sequences within *ssi* promoters and other repeat regions, also potential *ssi* elements (**Fig. 7A**). The homology of the ParB-like protein to ParB in a helix-turn-helix DNA binding domain (**Table S6**) indicates a regulatory function via sequence-specific DNA-binding. Our knockouts confirmed a role for the operon including this *parB*-like gene in transfer, consistent with the known role of the *ssi3* operon in establishment of immigrant plasmids (Golub *et al*., 1988; Al Mamun *et al*., 2021). Past work indicates genes in plasmid leading regions can interact with each other to determine transfer and stability, including those encoded by predicted *par* genes (Guynet *et al*., 2011). In a further experiment we found some evidence for such interactions: in a complementation assay, *in trans* expression of the ParB-like function in recipient cells increased transfer rate for wild-type plasmids, but not for knock-out plasmids (**Fig. S15**). In summary, our results implicate the *ssi3* operon not only as a target for plasmid evolution, but as a determinant of transfer rate, potentially via interactions among genes within the operon including the parB-like gene.

Another key implication of our results is that evolutionary stabilization of plasmids is host-specific (depends on the bacteria), demonstrated for example by the increased transfer rates we observed only with strain 15. The epistastic effects of mutations affecting transfer rate, which we observed upon deletion of the *ssi3* operon in p1ESBL or p15ESBL (**Fig. 7C**), help to explain this strain-specificity in the evolution of plasmids. One possible molecular driver of these epistatic effects, which could be explored further for other plasmids in future work, is interaction between resident and newly acquired plasmids. Our data are consistent with such interactions, in that strain 1 carries an IncF plasmid which also encodes the *ssi3* operon. This allows *in trans* expression of this operon from the resident plasmid, even in cells carrying the knocked-out version of p1ESBL or p15ESBL, and likely compensates for its deletion in the newly acquired ESBL-plasmids (**Fig. S14**). That expression of *psiB* in the *ssi3* operon is host-specific has previously been shown, suggesting this is also the case for other ORFs in this operon (Althorpe *et al*., 1999; Baharoglu *et al*., 2010). Thus, epistatic effects of mutations affecting plasmid stability traits may be common, and the likely role of plasmid interactions again emphasizes the importance of working with clinical/natural strains in order to identify the most relevant interactions.

In conclusion, our results show that rapid, strain-specific evolution of plasmid stability traits is a key predictor of which novel bacterium-plasmid combinations are most successful in the long term (over hundreds of generations and without antibiotic selection). Our findings address and go beyond the recent proposal to take compensatory evolution into account for such predictions (San Millán, 2018), by showing that rapid evolution of transfer rates as well as growth costs should be accounted for here. This points to some key pathways for improved prediction, and ultimately management, of problematic (highly stable and successful) plasmid-bacterium combinations. First, to further characterise the molecular signatures of successful combinations, such as the genetic changes and loci driving high transfer rates in our experiments, evolve-and-resequence experiments (Brockhurst *et al*., 2019) could be combined with analyses of phenotypic and genotypic changes for a wider range of strains and plasmids. Second, because many of the genetic changes driving our results were in loci specific to clinical strains (e.g., mobile elements, PAIs), there is a clear increase in the value of such data when they come from the most relevant clinical strains/plasmids, such as those presently circulating in patients and hospitals. Third, although plasmid stability traits measured in vitro can correlate well with in vivo properties (Benz *et al*., 2020), there are key differences such as spatial structure, the role of the host and resident microbiota. Therefore, downstream applications of candidate molecular signatures and rapid, strain-specific evolution as revealed by our experiments will benefit from validation in *in vivo* infection models.

## Material and Methods

### Bacterial strains and growth conditions

Strains 1, 15, and 19 were sampled from different patients in a transmission study at the University Hospital Basel, Switzerland. We have previously described phenotypes and genotypes of ancestral strains 1 (ST117, GCA_016433325.1) and 15 (ST40, GCA_008370755.1) and the genome sequence of the ancestral strain 19 (ST131, commonly associated with ESBL-plasmids, GCA_020907505.1) (Tschudin-Sutter *et al*., 2016; Benz *et al*., 2020; Huisman *et al*., 2021). These strains are natively associated with conjugative ESBL-plasmids of the type IncIγ, non-conjugative IncF for strain 19, and carry a range of other (non-focal, non-ESBL) plasmids (**Fig. 1**). Focal ESBL-plasmids all encode at least one TA system (pndA/B, and RHH-RelE for p1ESBL and pndA/B only for p15ESBL). Unless stated otherwise, we grew bacterial cultures at 37 °C and under agitation (180 rpm) in lysogenic broth (LB) medium, supplemented with appropriate amounts of antibiotics (none, 100 μg/mL Ampicillin (Amp), 50 μg/mL Kanamycin (Kan), 25 μg/mL Chloramphenicol (Cm)). We performed experiments with 4-6 biological replicates, with the exception of conjugation assays to estimate plasmid-transfer rates (n = 3). Culture volumes were 5 mL for culture tubes and 150μL for 96-well plates. We stored isolates in 25% glycerol at −80 °C.

### Construction of new strain-plasmid combinations

Generated strain-plasmid pairs are shown in **Fig. 1** (also see **Table S3**). To cure strains 1, 15 and 19 of their IncIγ-plasmids, we used replicon incompatibility and challenged each strain with the temperature-sensitive and Cm resistant pCP20-IncI1, encoding the replicon of p15ESL (Bakkeren *et al*., 2021). To cure p19ESBL, we used the Cm resistant pCure2 (Hale *et al*., 2010) competing with IncF replicons. We electroporated curing plasmids into the strains to be cured, plated them on the respective antibiotic for their overnight selection, and verified ESBL-plasmid loss by restreaking on selective plates and colony PCR in case of strain19-IncIγ. Curing plasmids were lost by growing ESBL-plasmid-cured colonies overnight at 37°C without antibiotic selection. We then combinatorially re-introduced focal ESBL plasmids into each *E. coli* host strain, by first conjugating strains 1 and 15 with the *Salmonella enterica* Typhimurium (S.Tm) strain ATCC 14028 *marT::cat* (Diard *et al*., 2017) to generate an intermediate host. Transferring p1ESBL and p15ESBL from S.Tm to the ESBL-plasmid cured recipients allowed the distinction of donor and recipient by plating on MacConkey agar (lac negative S.Tm forming yellow colonies and lac positive *E. coli* forming red colonies).

### Experimental Evolution

We serially passaged bacterial cultures (1:1000 dilution into 5mL fresh LB every day for 15 days; ∼150 generations in total) without antibiotics. To generate the starting populations, we grew ESBL-plasmid carrying and ESBL-plasmid-free strains from independent single colonies overnight and with the appropriate antibiotics. From these grown cultures, we transferred 2.5μL of ESBL-plasmid-carrying (p^+^) and of ESBL-plasmid-free (p^-^) cultures to their assigned tube. We mixed p^+^ and p^-^ cells at the start, rather than taking pure p^+^ cultures, so we could detect both net positive and net negative changes in frequency. On days 1, 3, 5, 10, and 15 we froze each bacterial population for long-term storage and plated them on LB-plates in dilutions appropriate for replicate-plating on Amp plates (quantitative detection limit of ESBL-carrying clones:plasmid-free clones at ∼1:100) and undiluted directly on Amp-plates to verify presence/absence of Amp resistant phenotypes. After 15 days, we additionally tested each replicate population for strain-genotype and ESBL-plasmid presence by PCR. All except for one resistant (and thus potentially p^+^) clone tested positive for the ESBL-plasmid by PCR (3 out of 72 tested colonies, see Supplementary Methods). We therefore take resistance-phenotype as a proxy for ESBL-plasmid presence. In addition to the four replicate populations per strain-plasmid pair, we passaged four ESBL-plasmid-free control populations for each strain. After 15 days of serial passage, we isolated two single clones, one p^+^ and one p^-^, from each endpoint population for phenotypic testing and whole-genome sequencing. One replicate population of strain1-p15ESBL (green) was contaminated (see Supplementary Methods) and thus excluded.

### Measuring plasmid-stability traits

To test for a growth cost associated with ESBL-plasmid carriage in the absence of antibiotics, we measured the population growth rates (*ψ*) of plasmid-carrying and corresponding plasmid-free clones in monocultures. We grew bacterial cultures in 96-well plates overnight, removed Amp from cultures of ESBL-plasmid-carrying clones by pelleting all cultures with centrifugation and subsequent resuspension, and transferred ∼1μL to a plate containing fresh LB without antibiotics with a pin replicator. These cultures grew for 24 h without agitation, and we estimated growth rates (h^−1^) based on eleven manual optical density (OD) measurements (Tecan NanoQuant Infinite M200 Pro) using the R package Growthcurver (Sprouffske and Wagner, 2016). To calculate the growth rates of p^+^ clones relative to p^-^ equivalents, we divided the growth rate of each p^+^ replicate by the averaged growth rate of the corresponding p^-^ clone.

To estimate ESBL-plasmid transfer rates *γ*, [mL (CFU h)^-1^]), we used the Approximate Extended Simonsen Model, accounting for varying growth rates of donor, recipient and transconjugants and estimating a time window for reliable estimation of *γ* (Huisman *et al*., 2022). ESBL-plasmids were always transferred to their ancestral plasmid-free equivalent marked with the Kan-resistance plasmid pBGS18 (Spratt *et al*., 1986; Benz *et al*., 2020). In brief, we grew independent overnight cultures of donor and recipient strains in the appropriate antibiotics, washed them by pelleting and resuspending and added ∼1 μL of 6.5-fold diluted donor and recipient cultures into 150 μL fresh LB with a pin replicator (total ∼1000-fold dilution). Mating populations grew for 6 h (**Fig. 3/ 7C**; 5 h for **Fig. S15**) without shaking, which was well within the estimated critical time window to avoid results being influenced by substantial ESBL-plasmid transfer from emerging transconjugants (Huisman *et al*., 2022). To enumerate the final cell densities, we plated the mating cultures at the end of the conjugation assays on selective LB-plates, with a detection limit of ∼ 20 CFU/mL, corresponding to a single transconjugant colony. For transfer rate estimates in **Fig. 4B** we used corresponding growth rate estimates from **Fig. 4A**. For the other transfer rate estimates (**Fig. 7C**; **Fig. S15**) we estimated growth rates by growing donor and recipient replicate populations as initially 1000-fold diluted, antibiotic-free monocultures in the plate reader for 24 h, with hourly OD measurements. Because donor and recipient strains were of the same type, i.e. strains 1, 15 or 19, we assumed transconjugants to grow at equal rates to the corresponding donor strains. Finally, we estimated *γ* with the R-package conjugator (Huisman *et al*., 2022). We performed the transfer assay with complementation of *parB*-like gene in recipient strains (**Fig. S15**) as described above with some alterations: independent overnight cultures grew with 0.2% glucose to repress promoter pBad. We washed them twice by pelleting and resuspending and subcultures them for 1.5 h in 2% L-arabinose to induce expression of the *parB* -like gene. We washed cultures to remove antibiotics and adjusted them to OD (600nm) = 0.5, and mixed equal volumes of donor and recipient cultures to initiate the transfer assay.

We tested for evidence of plasmid instability due to segregational loss by measuring the change in the frequency of plasmid-carrying cells during overnight culture in the absence of antibiotics, starting with pure cultures of plasmid-carrying strains. Prior to the assay, we grew ESBL-plasmid carrying strains as independent overnight cultures with Amp and washed them by pelleting and resuspending. Prior to their pin-replication into a 96-well plate containing fresh LB without antibiotics, we diluted and plated cultures on LB-plates (t = 0 h). After 24 h of growth, we plated grown cultures on LB-plates and replica-plated LB-plates (t = 0 h and t = 24 h) on Amp plates to estimate the fraction of plasmid carrying cells. We performed these assays for a subset of strain-plasmid pairs, including all ancestral combinations and evolved combinations that showed either low final frequencies or high among-replicate variation in the main experiment. Note that plasmid frequency in these assays can reflect a balance of segregational loss plus horizontal re-acquisition and variable growth rates. Nevertheless, any drop in plasmid frequency after starting with a pure p^+^ culture indicates individual cells can transition from p^+^ to p^-^.

### Population dynamics model

We used a deterministic mathematical model, based on the model described by Simonsen et al (Simonsen *et al*., 1990) and modified by Huisman et al (Huisman *et al*., 2022), to simulate plasmid dynamics in each strain-plasmid combination. More detailed information about the model can be found in the Supplementary Model Information. These models assume plasmids to spread by mass action kinetics (in our experiments bacterial populations grew in a well-mixed environment: liquid batch cultures with shaking). We modified the Extended Simonsen Model (Huisman *et al*., 2022) by tracking only the dynamics of plasmid-carrying (P1) and plasmid-free (P0) cells, rather than donors, recipients and transconjugants as in the original model. The differential equations (1-3) describe the dynamics for these subpopulations (dP0 and dP1) and nutrient availability (dC) over time:

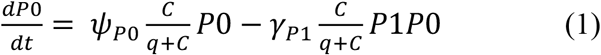

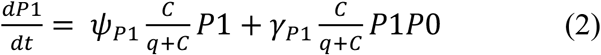

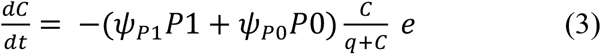

where cells grow at the maximal growth rate *ψ* (h^-1^) and plasmids are transferred at the maximal transfer rate *γ* (mL (CFU h)^-1^) from a plasmid-carrying to a plasmid-free bacterium. Both terms are regulated by the Monod function 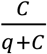, where C is the resource concentration (μg mL^-1^) and q the half-saturation constant (μg mL^-1^). E represents a conversion factor of resources into bacterial cells.

To simulate the expected change in plasmid frequency over time in each replicate and each strain-plasmid combination, we used experimentally determined values of *ψ* and *γ* for each ancestral (prior to experimental evolution) strain-plasmid pair, and for each replicate within each combination we estimated starting population densities based on plating (see Methods section Measuring plasmid stability traits and Supplementary Model Information section I). We compared plasmid dynamics in the models to those observed in our experiment by comparing simulated and observed values of the time-averaged plasmid frequency (area under the curve, AUC, for the time series of plasmid frequency over time for 15 days). We implemented the model in R (v4.1.0), using the function “ode” in the package deSolve for numerical integration (Soetaert *et al*., 2010).

We also used an expanded version of the model that accounts for evolutionary changes of plasmid-stability traits (Evo model; equations 4-10 in Supplementary Model Information section II). This model includes ancestral and evolved versions of bacteria (B_a_ and B_e_), with and without ancestral and evolved plasmids (P_a_ and P_e_). We simulated the dynamics of initially rare mutant strains, with mutations on the plasmid (B_a_P_e_), chromosome (B_e_ or B_e_P_a_) or both (B_e_P_e_), by including them in the model at an initial abundance of one cell at the start of the experiment. We empirically estimated *ψ* and *γ* for ancestral strains with and without ancestral plasmids (B_a_, B_a_P_a_) and evolved strains with and without evolved plasmids (B_e_, B_e_P_e_). We ran two versions of the model each making different assumptions about the unmeasured combinations B_a_P_e_ and B_e_P_a_ (see Supplementary Model Information section II).

### Sequencing and genomic analyses

Strains 1, 15 and 19 were previously sequenced with Illumina MiSeq (WGS, paired-end, 2 × 250 bp) and Oxford Nanopore MinION methods to generate hybrid assemblies (Huisman *et al*., 2021). Ancestral and evolved clones of the strain-plasmid combinations generated here (**Fig. 1**) were Illumina sequenced (NextSeq500, paired-end, 2 × 150 bp). In brief, we grew single clones in monoculture with appropriate antibiotics overnight and extracted genomic DNA (QIAGEN Genomic DNA Kit) for library preparation with the Nextera XT Library Preparation Kit. We used fastp for quality control, adapter trimming and quality filtering of reads acquired by Illumina sequencing (Chen *et al*., 2018). Because new strain-plasmid pairs could contain all plasmids from donor and recipient strains, we first generated the maximal reference sequence in silico against which we mapped new reads using breseq (v0.34.1) (Barrick *et al*., 2014). This revealed that after generating the new strain-plasmid combinations, all non-focal plasmids were retained in all strains. Further, we detected co-transfer of a non-focal plasmid (alongside the ESBL-plasmids) for a single non-focal plasmid (ColRNAI from strain 1 when generating strain19-p1ESBL). Based on this, we generated the final reference sequences used to compare ancestral and evolved genomes. We mapped reads for mutation calling with the breseq pipeline (v0.34.1) and used gdtools to correct mutations by corresponding ancestral sequences. Predicted mutations and unassigned missing coverage/ new junction evidence were curated manually for each sequenced clone using CLC Genomics Workbench 11 (Qiagen). To generate plasmid comparisons we used the visualization tool Easyfig (v2.2.2) (Sullivan *et al*., 2011). We visualized the genome and sequencing data using Circos (v0.69) (Krzywinski *et al*., 2009) and Geneious prime (v2022.1.1). For the ancestral ESBL-plasmid-free strain 19 sequencing resulted in poor quality and we mapped evolved ESBL-plasmid-free clones against the ancestral strain19-p1ESBL/ -p15ESBL instead. Sequencing revealed contamination in one replicate population of Strain1-p15ESBL, which we excluded from further analysis.

### Construction of plasmid mutants and expression vector

To test the effect of mutations in the *ssi3* operon of the plasmid leading regions, as we have observed them in both ESBL-plasmids with strain 15, we generated the two knockout mutants, mutA and mutB. For mutA we deleted the 2049bp large *parB*-like gene, thereby truncating the overlapping C-terminal of the hypothetical protein encoded by *ykfF* (34 nt). For mutB we deleted the *parB*-like gene and *psiA*/*psiB*, thereby truncating the genes *ykfF* and the N-terminal coding region of *ygaA*, encoding a hypothetical protein (removed 3 nt but retained a start codon). Likely the mutations exert polar effects with regard to downstream genes in this operon (**Fig. 7A** and **B**). To generate these mutants we used the lambda red recombinase system with pKD4 as a template for the Kan resistance marker (Datsenko and Wanner, 2000) (primers in **Table S11**). For the mutant construction, we conjugated p1ESBL and p15 ESBL to *E. coli* DH10B with pSIM18 (pSIM18 encodes λ Red recombination genes, facilitating homologous recombination, resistant to Hygromycin (Sharan *et al*., 2009)). We PCR verified Amp/Kan resistant colonies for insertion of the resistance marker. To remove the Kan cassette we introduced pFLP2, which encodes a FLP-recombinase and for which we replaced the Amp resistance marker with Cm (Hoang *et al*., 1998). For this, we grew strains containing the mutant plasmids at 37°C, leading to loss of the temperature-sensitive pSIM18 and allowing use of pSIM18 as a marker plasmid in strains 1/15/19ΔpESBL, for introduction of p1ESBLmutA/B and p15ESBLmutA/B via conjugation. We were not able to conjugate p1ESBLmutA/B and p15ESBLmutA/B into strain 19, which was also the case for the control-transferred WT p1ESBL/ p15ESBL in this experimental block. Therefore, we performed subsequent conjugation assays only with strains 1 and 15 (**Fig. 7C**). For the complementation assay, we cloned the *parB*-like gene from p1ESBL on the expression vector pHerd30T (Qiu *et al*., 2008), downstream of promoter pBAD employing USER cloning (New England Biolabs,(Cavaleiro *et al*., 2015)). In brief, we amplified the *parB*-like gene and pHerd30T by PCR with the Phusion U Hot Start DNA Polymerase (Thermo), employing primers with overhangs containing a Uracil base (primers p1ESBL; pFB135: acatacccAUGCCATCCGTGATTTCGGG/ pFB137: acggccagUtcaggcg-gcatcagccag, primers pHerd30T; pFB136: atgggtatgUatatctccttcttaaag/ pFB6 actggccgucg-ttttac, (5’>3’)).

### Statistics

We performed statistical analyses using R (v4.1.0). To test strain and plasmid effects for plasmid success (AUC in the main experiment) and plasmid stability traits (transfer rates and growth costs), we performed analyses of variances by fitting linear models (function ‘lm’). To calculate AUC, for both simulated and observed plasmid frequencies over time, we used the function “sintegral” in the package Bolstad2 for numerical integration (Bolstad, 2012). In analysing transfer rates, we assigned a dummy value of 10^−17^ (approximating the transfer rate if transconjugants would be present at the detection limit of ∼200CFU/mL) to the individual replicates where no transconjugants were detected (two replicates each for strain19-p1ESBL pink/green and the ancestral strain19-p1ESBL). Finally, to assess the fit of our different model variants, we performed linear regression of predicted and observed AUC. Note that for this we excluded the replicate population 3/ orange in Anc/ Evo Models, because of missing parameter estimates for this replicate caused by the lack of Amp-susceptible clones at day 15.

## Acknowledgements

We thank the Pathogen Ecology and Theoretical Biology groups at ETH Zürich, Martin Ackermann (ETH Zürich) and Álvaro San Millán (CSIC Madrid) for discussion and feedback. We thank Katia R. Pfrunder-Cardozo for help with the library preparation for sequencing, Niklaus Zemp from the Genetic Diversity Centre for help with the sequence analysis, Ricardo León-Sampedro for help with the genome plots and Adrian Egli from the from University Hospital Basel for providing bacterial strains.

## Supplementary Material

### 1) Supplementary Methods

#### Verification of plasmid carriage in evolved clones

In addition to sequencing of evolved (endpoint, after 15d) clones (colony isolates), we verified the presence of ESBL-plasmids by PCR-testing three further Amp-resistant clones from each endpoint population (**Table S1**). Because plasmid regions selected for PCR verification could in theory change during the evolution experiment, we used up to two distinct sets of primers for each clone. We amplifed the *oriT* region of both IncIγ plasmids (pIncI_oriT_fw: AGTTCCTCATCGGTCATGTC/ pIncI_oriT_rv: GAAGCCATTGGCACTTTCTC) and used one primer pair specific to p1ESBL (p1ESBL_fw: GGAACCCGAAGTATCTCACC/ p1ESBL_rv: GGAGCACCA-GAAAGTCCAC) or p15ESBL (p15ESBL_fw: AGGATGA-ACTGAAGGCAACG/ p15ESBL_rv: CCAGTAGATAAACCACATTGC). This allowed us to verify plasmid presence with at least one of the primer pairs for three clones of 22 of the 23 endpoint populations. For one population of strain 15-p15ESBL (purple), we could not confirm plasmid carriage in the three tested clones with these primers, although the sequenced clone from this population did carry the plasmid.

#### Verification of strain identity in endpoint populations

We verified strain identity for multiple colonies from each endpoint population. Because strains 1, 15 and 19 belong to different sequencing types and phylogroups we could differentiate them by Clermont phylo-typing (**Table S2**) (Clermont *et al*., 2000, 2013). For strain 1, this resulted in amplification of *chuA* (288bp, indicating group F), for strain 15 in amplification of *ArpA* (400bp, indicating B1), and for strain 19 *chuA* and *yjaA* (211bp, indicating group B2). For each endpoint population, we tested three colonies, three per colony morphology in case of multiple distinct colony morphologies, and repeated the evolution experiment for replicate populations where contamination was suspected. For the strain1-p15ESBL_green population, we noticed the contamination only later and therefore this population was excluded entirely. We verified that plasmid-free control populations remained susceptible to Amp throughout the whole experiment by selective plating of undiluted cultures. In cases where we found growth, we concluded cross-contamination with plasmid-containing populations and repeated the evolution experiment for those replicates.

### 2) Supplementary Model Information

We used a mathematical model with two main variations variations: the ancestral (Anc) model and the evolved (Evo) model, each with two different starting conditions.

#### I) The ancestral model

To simulate plasmid dynamics during the 15-day serial passage, assuming plasmid-carrying and plasmid-free subpopulations maintain their ancestral phenotypes (the phenotypes of the different ancestral strain-plasmid combinations and plasmid-free equivalents prior to the 15-day experiment), we used existing models (Simonsen *et al*., 1990; Huisman *et al*., 2022) with a simplified notation. Because the donor and recipient strains in each of our replicate populations were otherwise identical (to the best of our knowledge), we assumed transconjugants to be identical to donors. This allowed us to simplify the notation from existing models (with subtypes donors, recipients and transconjugants) to have only two subtypes (plasmid carrying, P1, and plasmid-free, P0; **Fig. S4**). Note this change in the notation simplifies the equations and plotting, but does not affect the results (if we instead use the original models but sum the abundances of donors and transconjugants together, we obtain the same output). In these models, each subtype replicates at their specific growth rate *ψ*, constrained by nutrient-limitation (Monod function), and plasmids can transfer at rate *γ* to plasmid-free cells. We estimated both parameters for each strain-plasmid pair experimentally (**Fig. 3A-B**), and estimated starting densities of P0 and P1 (*t = 0*) for each replicate population by plating. To simulate serial passage, each 15-day simulation included 15 sequential growth cycles, with each growth cycle except the first one initiated with 0.001 times the abundances from the preceding cycle. To initiate the first growth cycle, we used abundances estimated by plating.

#### II) The evolved model

In the evolved model (Evo Model) all six combinations of host (ancestral, B_a_, and evolved, B_e_, bacteria) and plasmid (ancestral, P_a_, and evolved, P_e_) were permitted (**Fig. S5**). We could only measure the plasmid stability traits *ψ* and *γ* for either fully ancestral or fully evolved pairs. We therefore made two versions of the Evo Model, each making different assumptions for the growth rates of B_e_P_a_ and B_a_P_e_. (i) In Evo Model 1, the evolutionary state of the plasmid decides the growth rate. Therefore 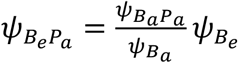 and 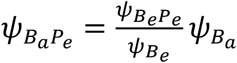. That is, B_e_P_a_ grows like B_e_, but pays the same cost of carrying P_a_ as B_a_ does; likewise, B_a_P_e_ grows like B_a_, but pays the same cost of carrying P_e_ as for B_e_ does. (ii) In Evo Model 2, the evolutionary state of the bacterium decides the growth cost of plasmid carriage for combinations not measured directly. Therefore, in this version of the model, 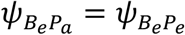 and 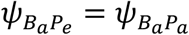. For *γ* of the subtypes B_e_P_a_ and B_a_P_e_, we assumed this was determined by the evolutionary state of the plasmid and therefore 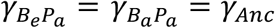 and 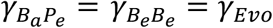, and we applied this assumption for both Evo Models, 1 and 2. For single (evolved plasmid or bacterium) and double (evolved bacterium with evolved plasmid) mutants, we assumed invasion from a very low frequency by taking the initial abundance of each mutant type to be one cell (at *t* = 0). For other initial population densities we followed the same rationale as for the Anc Model.

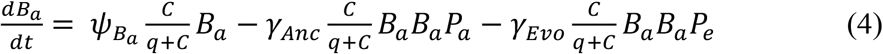

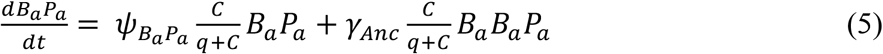

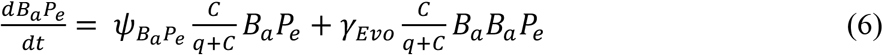

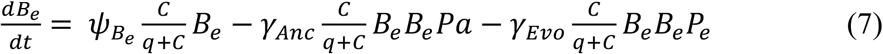

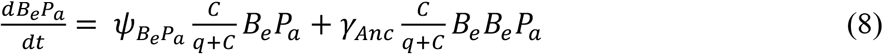

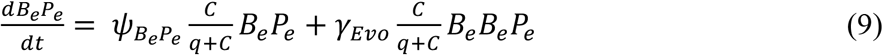

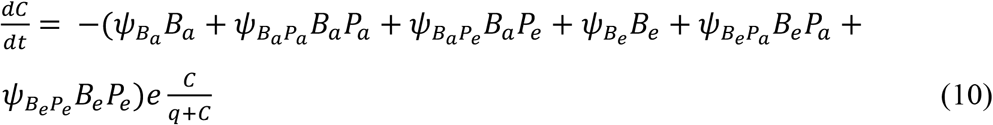

#### III) Resource availability and the resulting population densities

For all the model variations above, we set the amount of resources required per cell division to *e* = 1, which allows for maximal population densities identical to the initial resource concentration C_0_. For the half-velocity constant, we assumed *q* = 0.5 × *C*_0_. For Anc and Evo Models we set C_0_ = 1.15 × 10^12^ and therefore number of cells produced per growth cycle *N* = 1.15 × 10^12^. This setting predicted plasmid dynamics close to the observed plasmid frequencies, for example with some plasmids spreading and others declining in frequency, while overestimating total population densities. An alternative model, where we estimated C_0_ by calculating the average of total population density enumerated on days 1,3,5,10, and 15 (*N* = 1.15 × 10^9^ = *C*_0_, **Fig. S6**), yielded reasonable total population sizes, but consistently underestimated plasmid frequencies. For example, in this version of the Anc model all plasmids were eventually lost (albeit at variable rates), unlike in our 15-day experiment. Despite this, the predictions arising from these two versions of the model (with larger and smaller total population sizes, due to different values of C_0_) were similar (correlation of simulated AUCs: *r* = 0.7, *p* < 0.001 for Anc Model; *r* = 0.48, *p* < 0.05 for Evo Model 1; *r* = 0.42, *p* < 0.05 for Evo Model 2 comparison). For example, both versions of the Evo Model 1 captured spread of strain 19-p1ESBL better than both ancestral versions. Note that these models, as in previous work with the same basic model (Simonsen *et al*., 1990), assume no transfer in stationary phase. This assumption potentially contributes to the lower-than-observed average plasmid frequencies observed with realistic total population sizes (because if there is in fact some stationary phase transfer, the resulting transconjugants are not accounted for by such a model). We describe an alternative version of the model below that relaxes this assumption, allowing transfer in stationary phase.

#### IV) Testing the role of model complexity

Because simpler models contain fewer parameters, we may generally expect them to explain less variation than complex models with more variables. To test whether the higher predictive power of our Evo Models could be explained by their greater complexity alone (rather than the evolved parameters themselves being better predictors than ancestral parameters), we introduced the evolved parameters (measured for evolved p^+^ and p^-^ clones from each replicate population) into the ancestral model. This version of the model explained the experimentally observed variation of plasmid success as well as Evo Model 1 did, and better than Evo Model 2 did (*R*^*2*^ = 0.31, *p* < 0.05, **Fig. S7**). This shows the evolved phenotypes were responsible for the better fit of Evo Model 1 and 2 compared to the Anc Model, rather than the increased complexity (greater number of subtypes) of these versions of the model.

#### V) *Relaxing stationary growth phase assumptions for* γ

As in earlier work using the same models, the Monod function 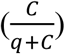 in the transfer-term means there is no plasmid transfer when there is no growth. In other words, it assumes there is no stationary phase plasmid transfer. For IncF-family plasmids, transfer is known to be reduced when bacterial growth is inhibited (Frost and Manchak, 1998; Headd and Bradford, 2020), suggesting this assumption is at least sometimes reasonable. However, how this translates to other plasmid types and growth conditions is not well understood. Therefore, the transfer assumption may be overly stringent in some cases, and a possible explanation for the underestimation of average plasmid frequency in our models above that otherwise reproduced total population growth relatively realistically. To test this, we detached plasmid transfer from resource availability in the Anc Model and Evo Model 1. We did this by removing 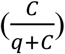 from the transfer terms (terms including *γ* (Anc Model; eq. 1-2) or *γ*_*Anc*_ or *γ*_*Evo*_ (Evo Model; eq. 4-9)) in each equation. With realistic total population sizes (*C*_0_ = 1.15 × 10^9^ = *N*), the predictions of this version of the model were identical to the stricter versions above which assumed no stationary phase transfer (correlation between simulated AUC values for the two model versions: *r* = 1 for both the Anc model and Evo Model 1). With *C*_0_ = 1.15 × 10^12^(resulting in more total population growth), the model allowing stationary phase transfer differed from the more stringent version and performed relatively well (correlation between observed and simulated AUC values: *R*^*2*^ = 0.66, *p* < 0.001 for Anc Model; *R*^*2*^ = 0.64 *p* < 0.001 for Evo Model 1; **Fig. S8**). Despite the good performance of this model, it should be interpreted with caution, because it assumes both unrealistic total population growth and unrestrained stationary phase transfer. Nevertheless, this suggests future work with variations of these widely used population dynamic models should consider the possibility that this assumption may result in underestimation of plasmid transfer across the whole growth cycle.

### 3) Supplementary Figures

**Figure S1:**
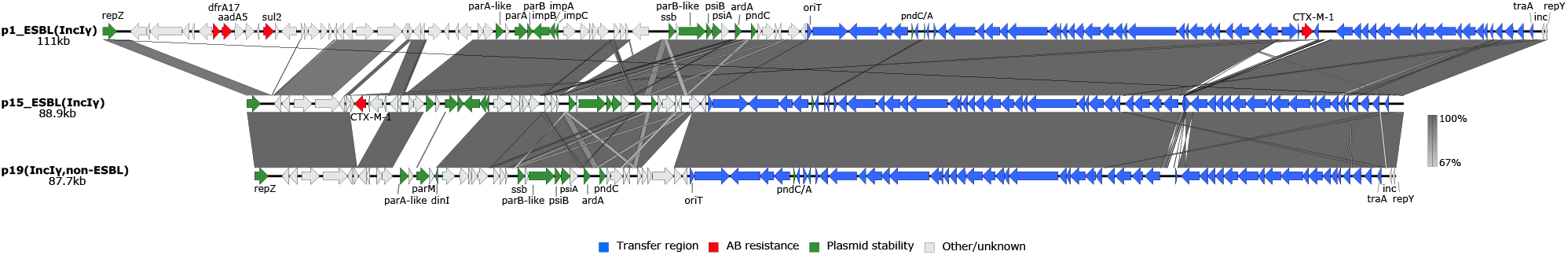
Comparison of IncIγ plasmids natively associated with the three clinical strains used in this study. All three strains (1, 15, 19) were natively associated with IncIγ plasmids (sequences shown here). The IncIγ plasmids from strains 1 and 15 (p1_ESBL and p15_ESBL; the top two sequences in this plot) encode ESBL phenotypes and were included in our experiments. The IncIγ plasmid from strain 19 (strain19_IncIγ; the bottom sequence in this plot) does not encode an ESBL phenotype and was therefore not included in our experiments, but is shown here to provide background information about the plasmids native to each strain. The three plasmids show high sequence similarity in their backbone. Gene functions are given in color (see legend at bottom), and for simplicity only selected genes are labelled.

**Figure S2:**
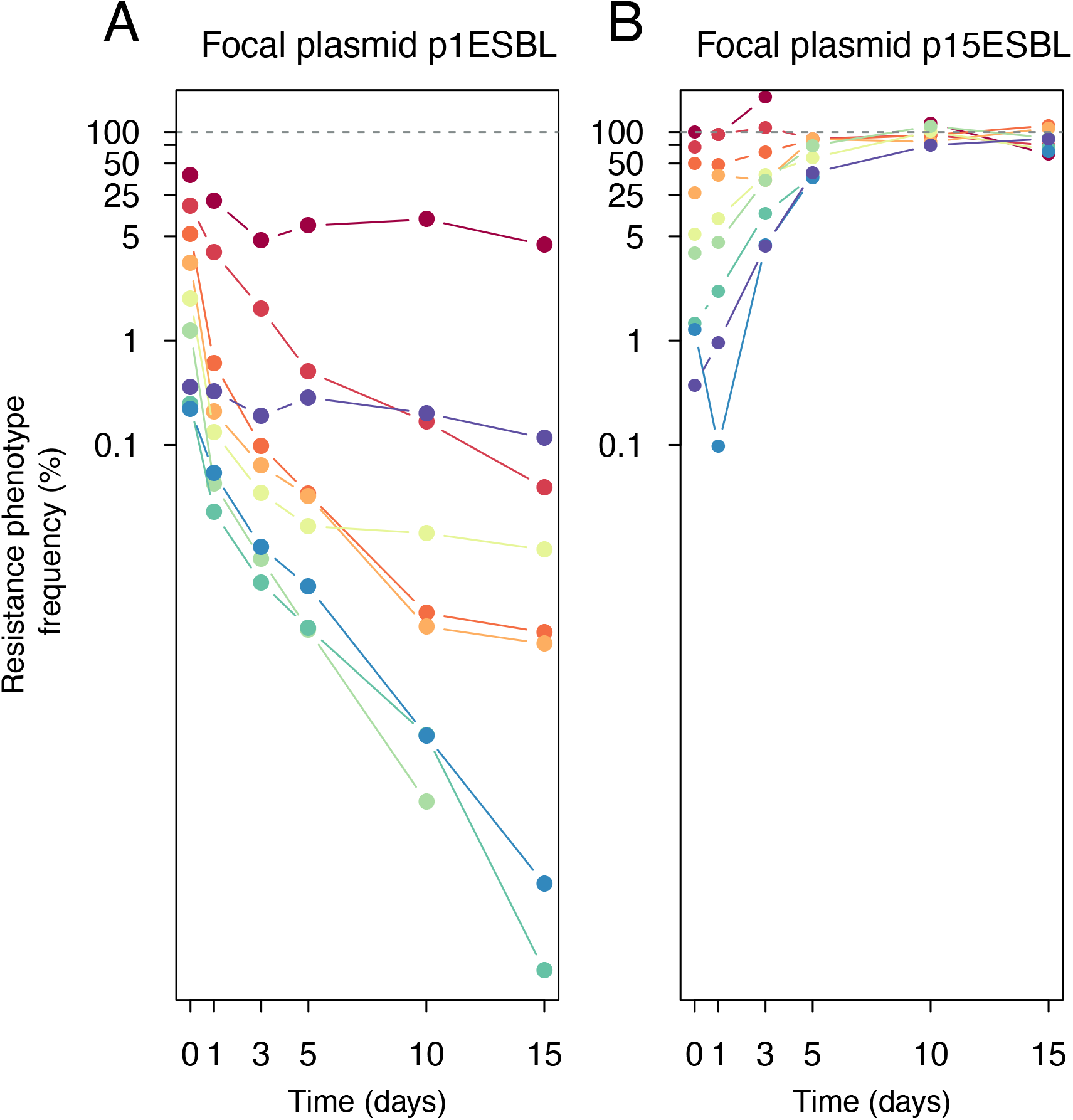
Testing whether initial plasmid frequency affects plasmid dynamics for strain 19 with p1ESBL (A) and p15ESBL (B) during an additional 15-day experiment. The key result here is that across a range of starting frequencies, p1ESBL declined during this 15 day experiment and p15ESBL increased in frequency. This shows any average difference in starting frequency in the main experiment is unlikely to have explained the difference in plasmid success in **Fig. 2**. We performed this additional evolution experiment in the same way as that described in the main text (**Fig. 2**; Methods), with a slightly different plating scheme to estimate the fraction of resistant phenotypes (separate plating on LB and Amp agar, rather than replicate-plating). For Strain19-p1ESBL, we detected significant plasmid loss already after overnight culture in the presence of Amp prior to inoculation. This is in line with our other, short-term plasmid stability assays (**Fig. S3/S9**). For both strain-plasmid pairs (A&B), we varied the starting plasmid fraction by varying the ratios of pure cultures made from plasmid-carrying and plasmid-free strains mixed together at the start.

**Figure S3:**
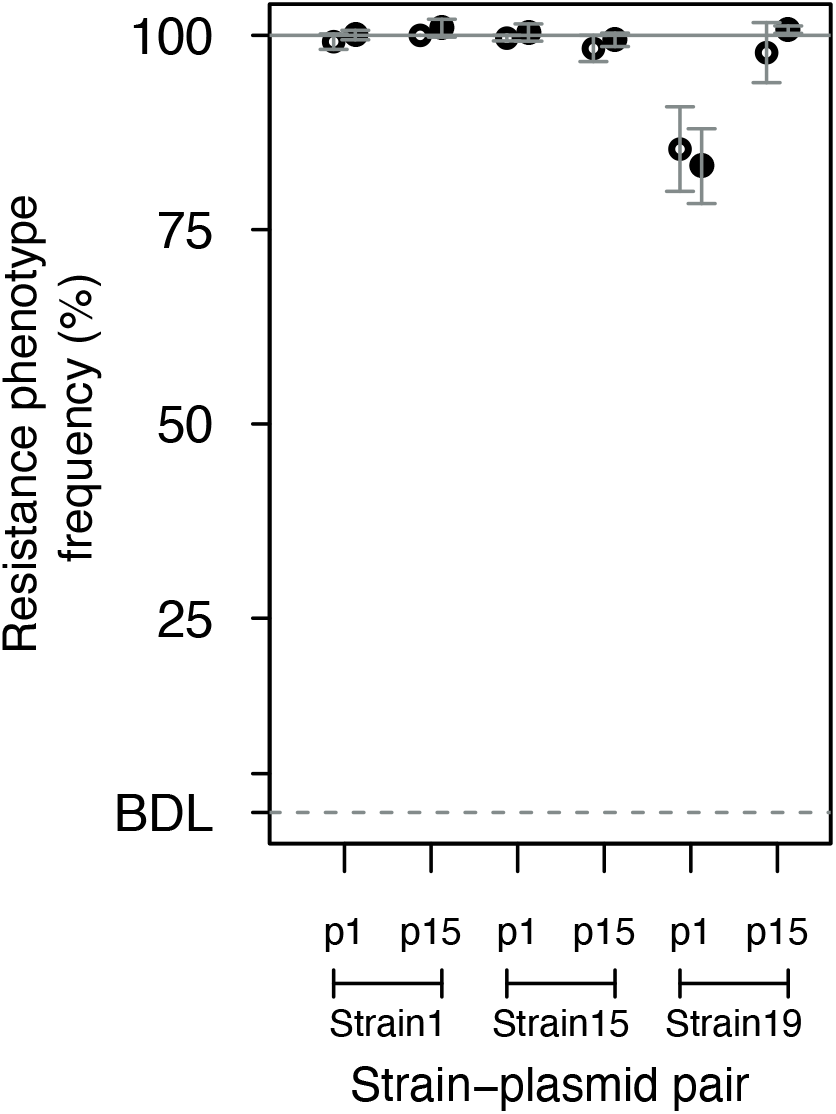
Stability of plasmids in each ancestral strain-plasmid combination over 24 hours. The frequency of plasmid-carrying cells was estimated by selective plating. For each strain-plasmid combination, empty circles show the fraction of plasmid-carrying cells at *t* = 0 h and full circles at *t* = 24 h. Each culture was inoculated from a pure-culture of a plasmid-carrying strain, so any net loss reflects significant segregational loss (relative to possible re-acquisition via conjugation in the same cultures; see Methods). The relatively low initial frequency for strain19-p1ESBL is consistent with our other observations (**Fig. S2/ S9**), suggesting possible plasmid loss during cultivation prior to inoculation. Despite this, this plasmid was stable after inoculation into the experimental conditions here (similar frequency at *t* = 0 h and at *t* = 24 h). Stability (the change in frequency over 24 h) was not dependent on strain or plasmid (two-way ANOVA with frequency at *t* = 24 h relative to frequency at *t* = 0 h as the response variable, effect of plasmid: *F*_1,25_ = 0.355, *p* > 0.05; effect of strain: *F*_2,25_ = 0.011, *p* > 0.05).

**Figure S4:**
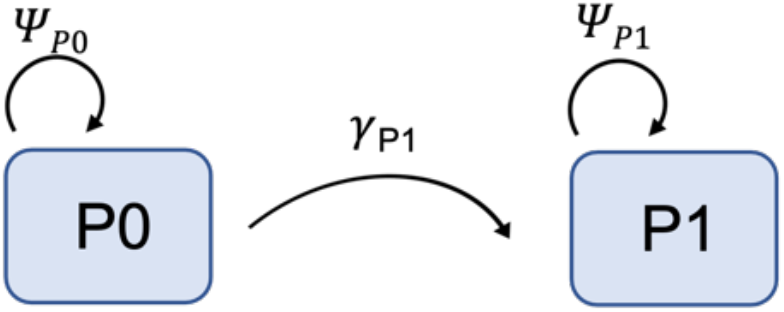
Schematic diagram of the Ancestral (Anc) model for plasmid population dynamics (equations 1-3). P0 denotes plasmid-free cells and P1 plasmid-carrying cells. Arrows indicate transition of cells between states, where *ψ* is bacterial replication and *γ* is horizontal plasmid transfer.

**Figure S5:**
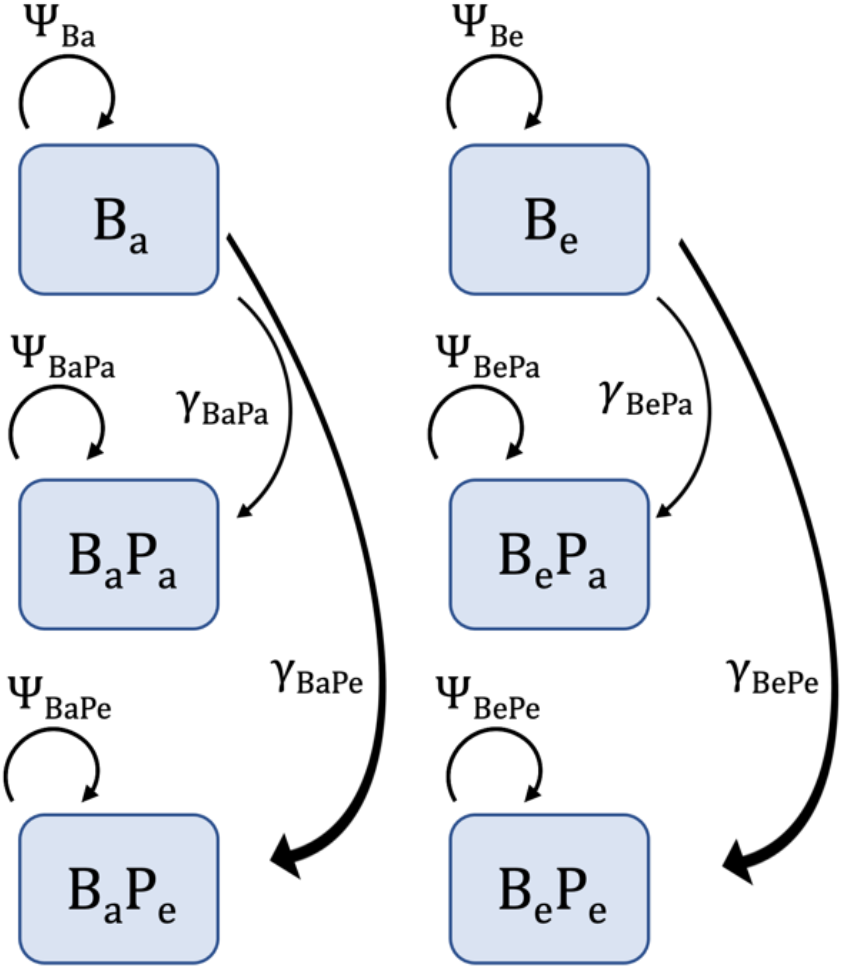
Schematic diagram of the Evolved (Evo) Model for plasmid population dynamics, which includes ancestral and evolved strain-plasmid pairs (equations 4-10). B_a_/B_e_ denote ancestral/evolved bacterial hosts. P_a_/P_e_ denote ancestral/evolved plasmids. Arrows indicate transition of cells between states, as above in **Fig. S4** and as described in the Supplementary Model Information section.

**Figure S6:**
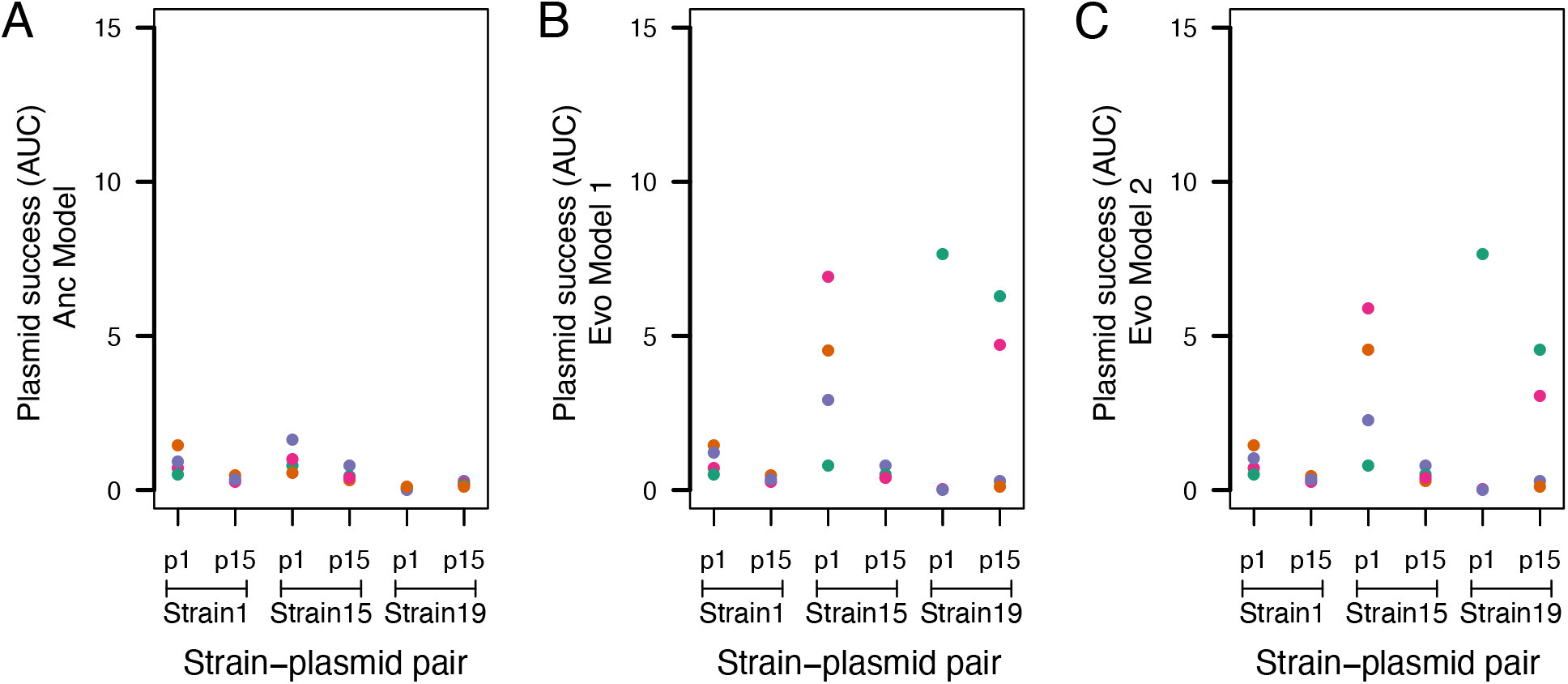
Plasmid success over time (area under the curve, AUC, from plasmid frequency over 15 days) for each replicate population in each strain-plasmid combination (x-axis), from simulations with alternative versions of each model. Here the total population size is smaller than in the versions shown in the main text (because here we set *C*_0_ = 1.15 × 10^9^). (**A)** Anc Model, (**B)** Evo Model 1, (**C)** Evo Model 2. Colors show replicate populations.

**Figure S7:**
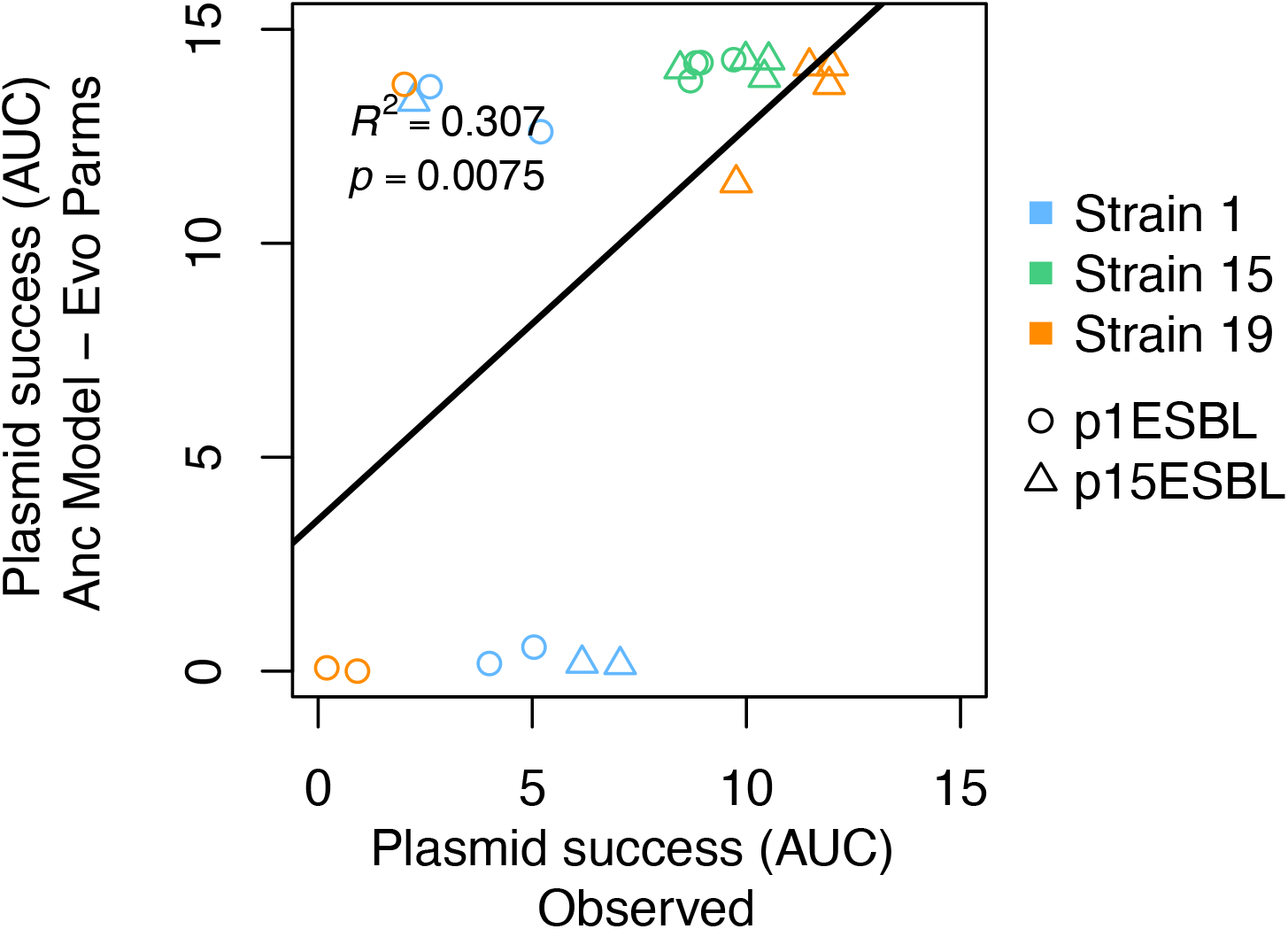
Observed plasmid success over 15 days vs predictions from the Anc Model implemented with evolved parameters. Plasmid success (area under the curve, AUC, from plasmid frequency over time) observed in our 15-day serial passage experiment (x-axis) is plotted against values predicted by the ancestral model, but implemented using evolved parameters (growth rates and transfer rates measured for evolved clones isolated at the end of the experiment; y-axis).

**Figure S8:**
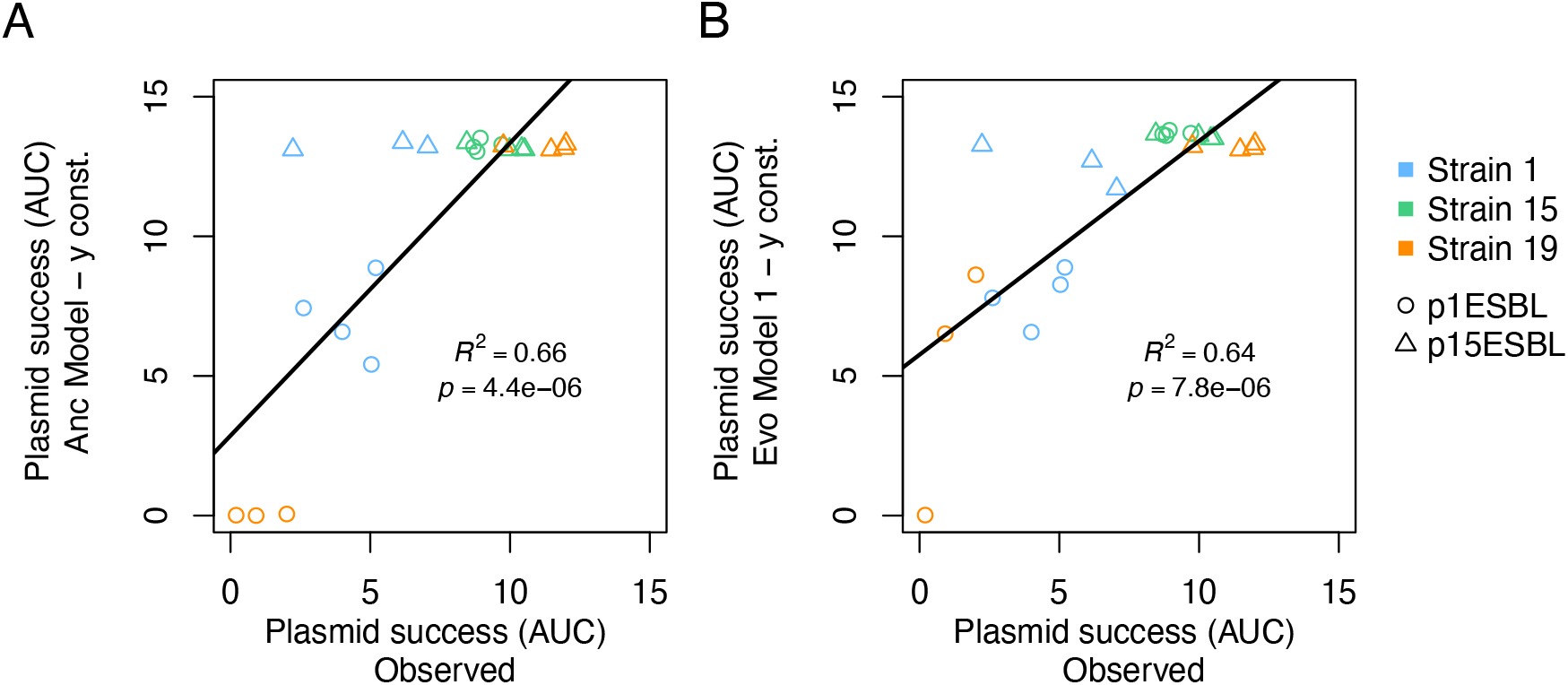
Observed plasmid success over 15 days vs predictions from (**A**) the Anc Model and (**B**) the Evo Model 1, but allowing for plasmid transfer in stationary phase. Plasmid success (area under the curve, AUC, from plasmid frequency over time) observed in our 15-day serial passage experiment (x-axis) is plotted against values predicted by alternative versions of the Anc and Evo Models (A&B), where the assumption that plasmid transfer is coupled to bacterial growth is relaxed (models shown in the main text, like previous work with the same models, assumes transfer is dependent on growth, via resource concentration). Note that these models assume both unrealistic total population growth (*C*_0_ = 1.15 × 10^1^) and unrestrained plasmid transfer, and should therefore be interpreted very cautiously. These assumptions explain the higher average plasmid frequencies (higher AUC) for successful combinations here compared to other versions. This in turn helps to explain the similar performance of Anc and Evo versions here (successful combinations already spread very rapidly in the Anc Model (top-right in panel A), so evolutionary changes increasing plasmid transfer rates or reducing growth costs that are incorporated in the Evo Model have little impact on predicted dynamics).

**Figure S9:**
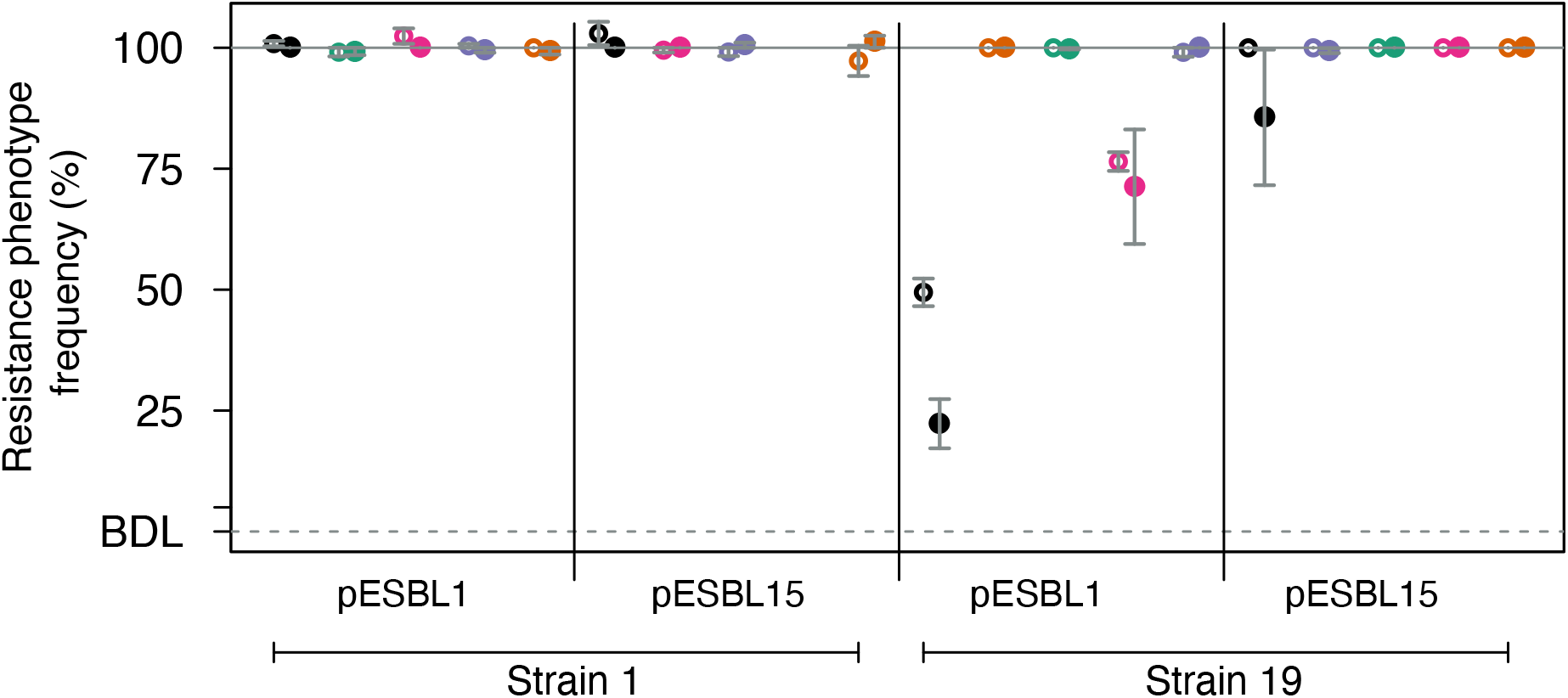
Plasmid stability over 24 hours for ancestral and evolved clones. The frequency of resistant colonies (approximating plasmid frequency) was estimated by plating at the start and end of 24 h incubation without antibiotics for strains 1 (left half) and 19 (right half) with pESBL1 and pESBL15. Black points indicate the ancestral clone in each combination; other colours indicate evolved clones, with different colours indicating which replicate population each evolved clone was isolated from (as in other figures). For each clone (each pair of adjacent points), empty and full circles show the fraction of plasmid-carrying cells at *t* = 0 h and *t* = 24 h, respectively. The ancestral clone of strain 19 (black) was the only clone showing clear loss of plasmids during the assay (from ∼50% to ∼20% for p1ESBL, and from ∼100% to ∼80% for p15ESBL). This is consistent with relatively low frequencies upon inoculation observed in other experiments (**Fig. S2 & S3**). However, among evolved clones all plasmids were highly stable. There was no evidence that stability (taken as frequency at 24 h relative to at 0 h for each replicate assay) varied among the different evolved clones within each strain-plasmid combination (one-way ANOVA within each of the four strain-plasmid combinations: *p* > 0.05 in all cases), or on average among the different combinations (two-way ANOVA with stability as the response variable, taking the average for each evolved clone: effect of plasmid: *F*_1,12_ = 3.414, *p* > 0.05; effect of strain: *F*_1,12_ = 1.405, *p* > 0.05).

**Figure S10:**
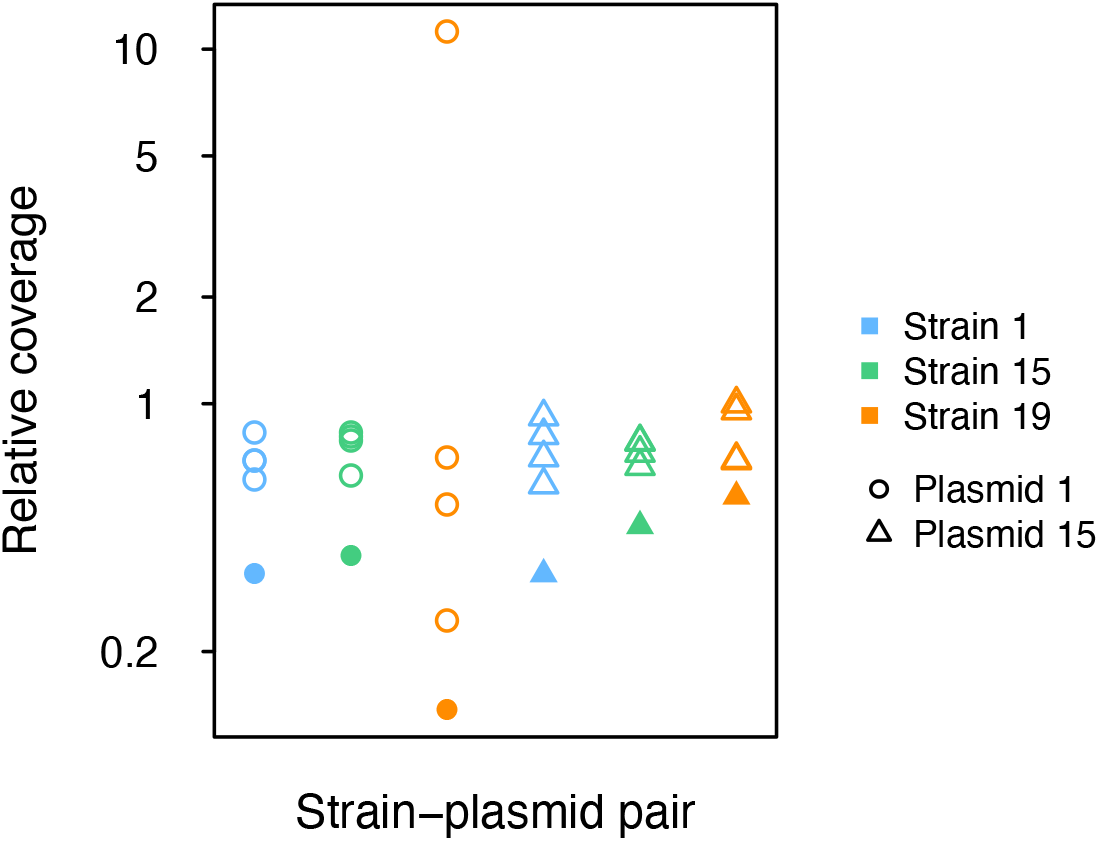
Coverage for ESBL-plasmid sequences relative to corresponding chromosomal sequences. Relative coverage (y-axis) is calculated as mean ESBL-plasmid coverage relative to mean chromosomal coverage for each clone in each strain-plasmid combination (see legend), with mean read-depth coverage estimated by breseq (Barrick *et al*., 2014). Evolved clones (empty symbols) consistently had higher ESBL-plasmid relative coverage compared to ancestral clones (6 out of 6 cases; filled symbols). In all cases except one, the difference was moderate. These moderate differences should be interpreted with caution, because sequence preparation and coverage estimation may be subject to different biases for plasmids and chromosomes. However one evolved clone of strain 19-plasmid 1 showed a much larger difference in relative coverage compared to the ancestor. For this evolved clone, this suggests an increased copy number of p1ESBL, which coincided with an increase in transfer rate of several orders of magnitude (**Fig. 4B**, evolved replicate 3/purple). These findings are also in line with a recent study demonstrating increased plasmid transfer rates after evolutionary increases in copy number (Dimitriu *et al*., 2021).

**Figure S11:**
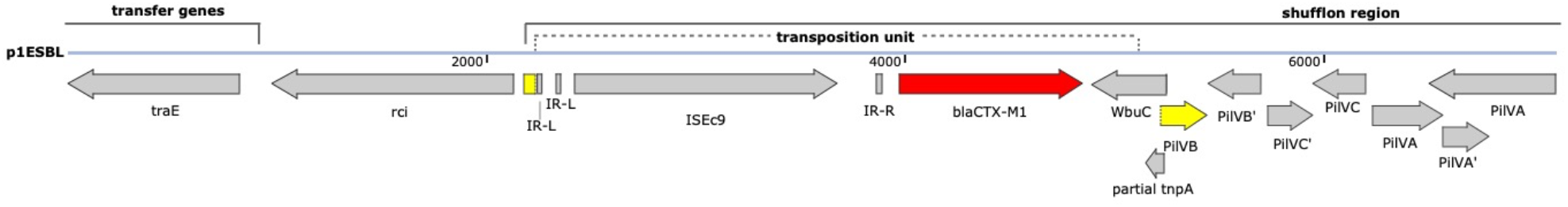
Shufflon region of the ancestral p1ESBL plasmid (position 88’151-96’017). A transposition unit containing an ISEc9 and *bla*_CTX-M1_ (red) disrupts pilVB (yellow) within the shufflon segment B. In one evolved clone from the end of our 15-day serial passage experiment (**Fig. 2**; strain 19-p1ESBL replicate 2), this transposition unit had jumped into the chromosome (in *astA*, aerobic arginine catabolism; **Fig 6C**, orange), and the plasmid itself had been lost. In two other evolved clones (strain 19-p1ESBL replicate 4, pink in **Fig. 6C** and strain 1-p1ESBL replicate 3, purple in **Fig. 6A**), the entire transposition unit was inverted.

**Figure S12:**
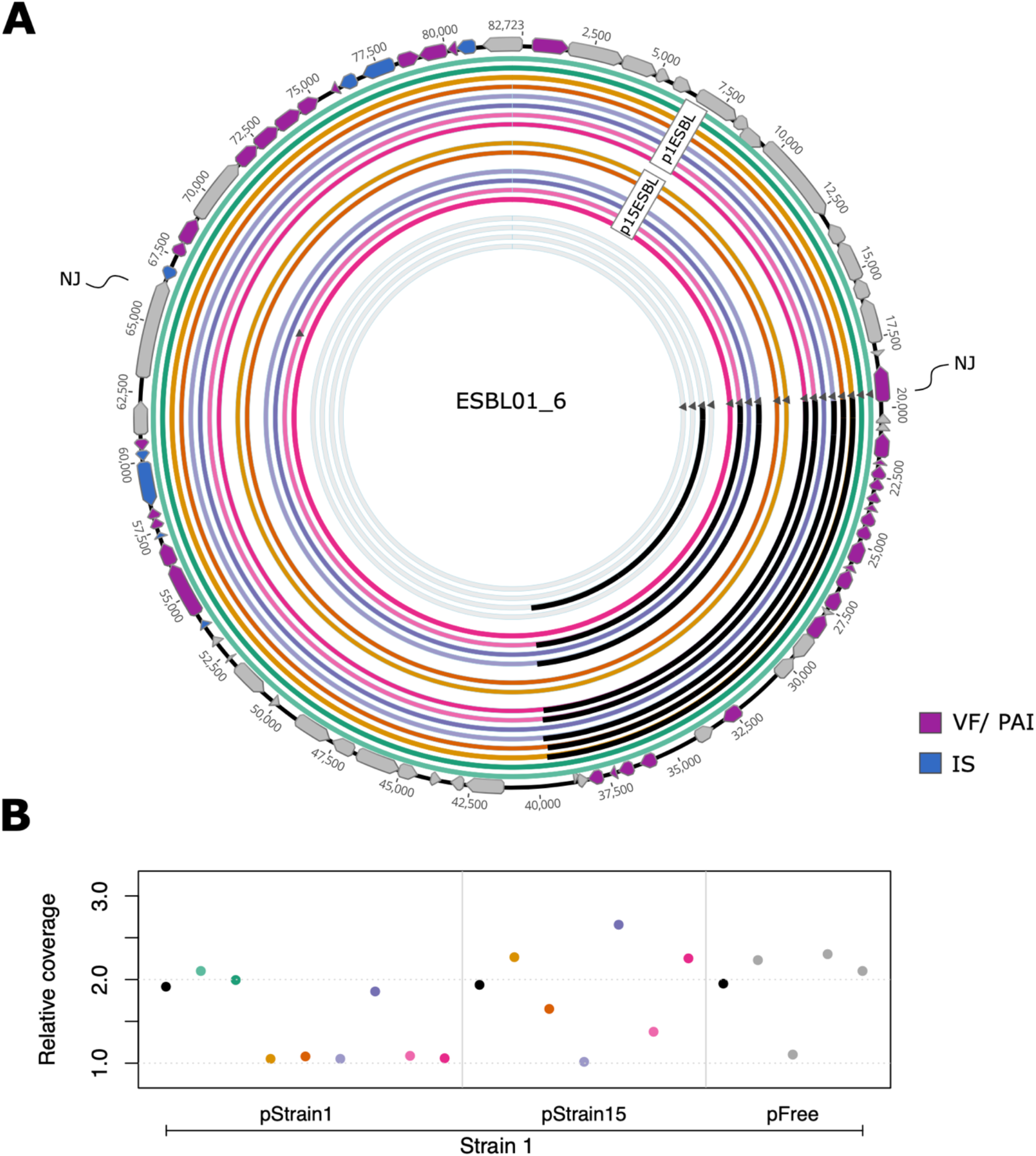
Deletions in a putative circularized pathogenicity island (PAI) in strain 1. **A)** With long-read sequencing we have previously determined the contig ESBL01_6 to be circularized in the original clinical isolate (Benz *et al*., 2020). Resequencing (Illumina) of the ancestral and evolved clones used in this study revealed a NJ (triangle) with the end of ESBL01_2, one of the two chromosomal contigs. This indicates that ESBL01_6 can exist both as part of the chromosome, potentially closing the gap between the two chromosomal contigs ESBL01_1 and ESBL01_2, and in circularized form. Black bars indicate ∼20kb deletions and the colour of annotated genes indicates their classification determined by VrProfile (purple = virulence factor and/ or pathogenicity island, blue = insertion sequence, **Table S8**) (Li *et al*., 2018). **B)** Sequence coverage (Illumina) of ESBL01_6 relative to the average of the other chromosomal contigs indicates ESBL01_6 to be present either as a single or two copies. This coincides with deletions in panel A: two copies are present in the absence of the deletion and only one copy is present in case of a deletion in the PAI.

**Figure S13:**
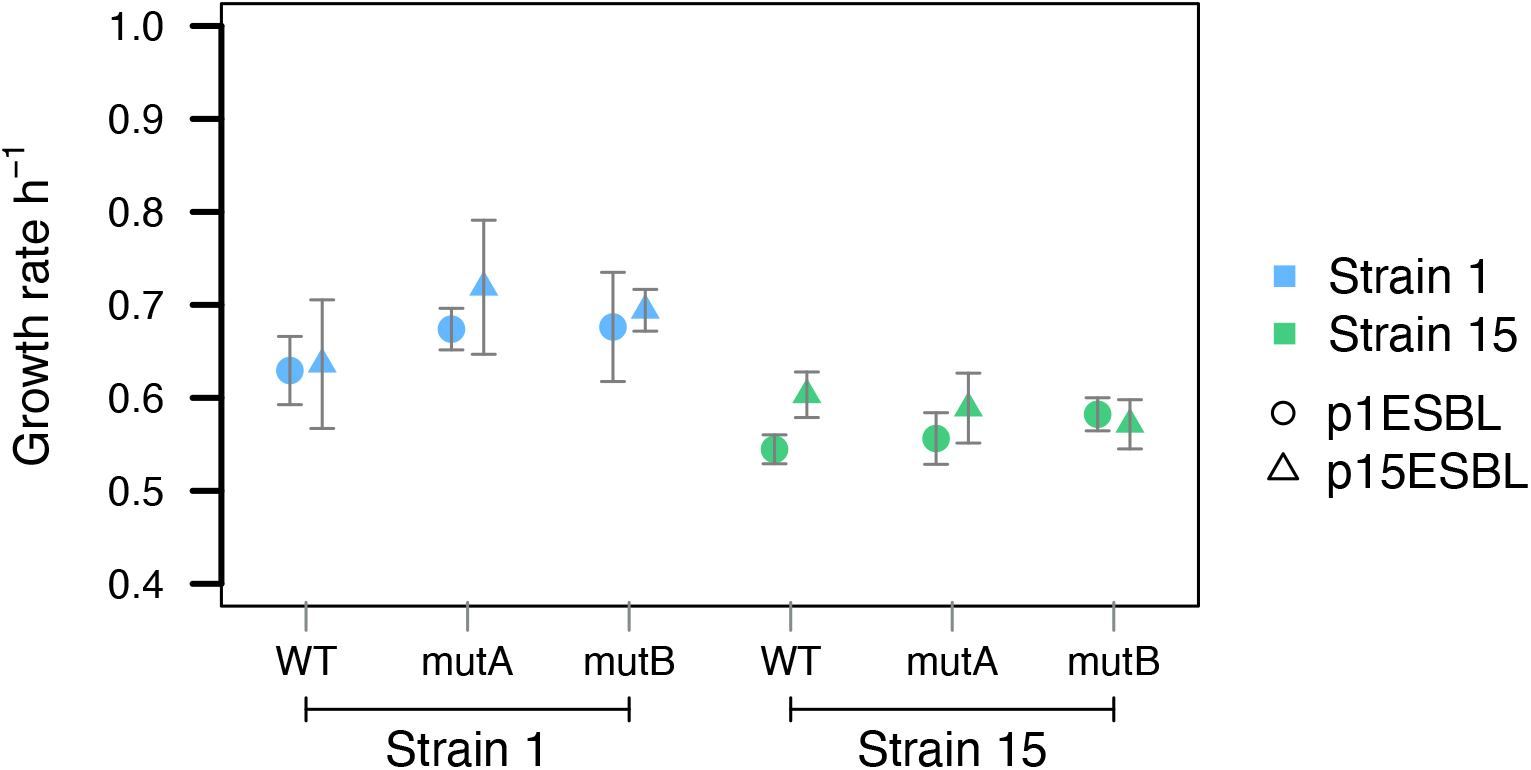
Generated leading-region mutations (mutA, mutB) have no significant effect on bacterial growth rate relative to the wild type (WT), measured in the absence of antibiotics over 24h (Welch Two Sample *t*-test, *p* < 0.05 in all cases before and after Holm’s correction for multiple testing).

**Figure S14:**
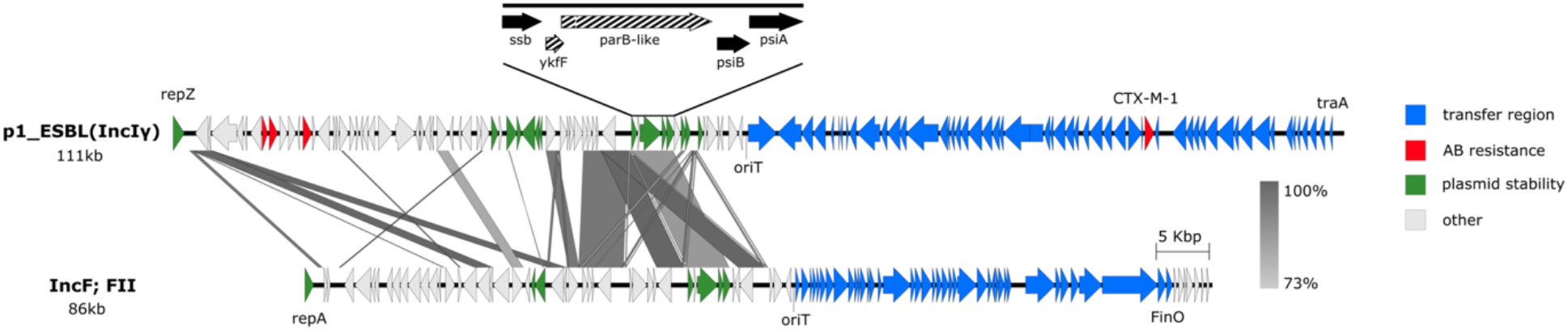
High sequence similarity of the *ssi3* operon on p1ESBL (shown here; top) and p15ESBL (not shown) with the *ssi3* operon on the IncF plasmid native to strain 1 (bottom). The *ssi3* operon has been shown to be largely conserved among conjugative plasmids in *E. coli* (Wein et al., 2021).

**Figure S15:**
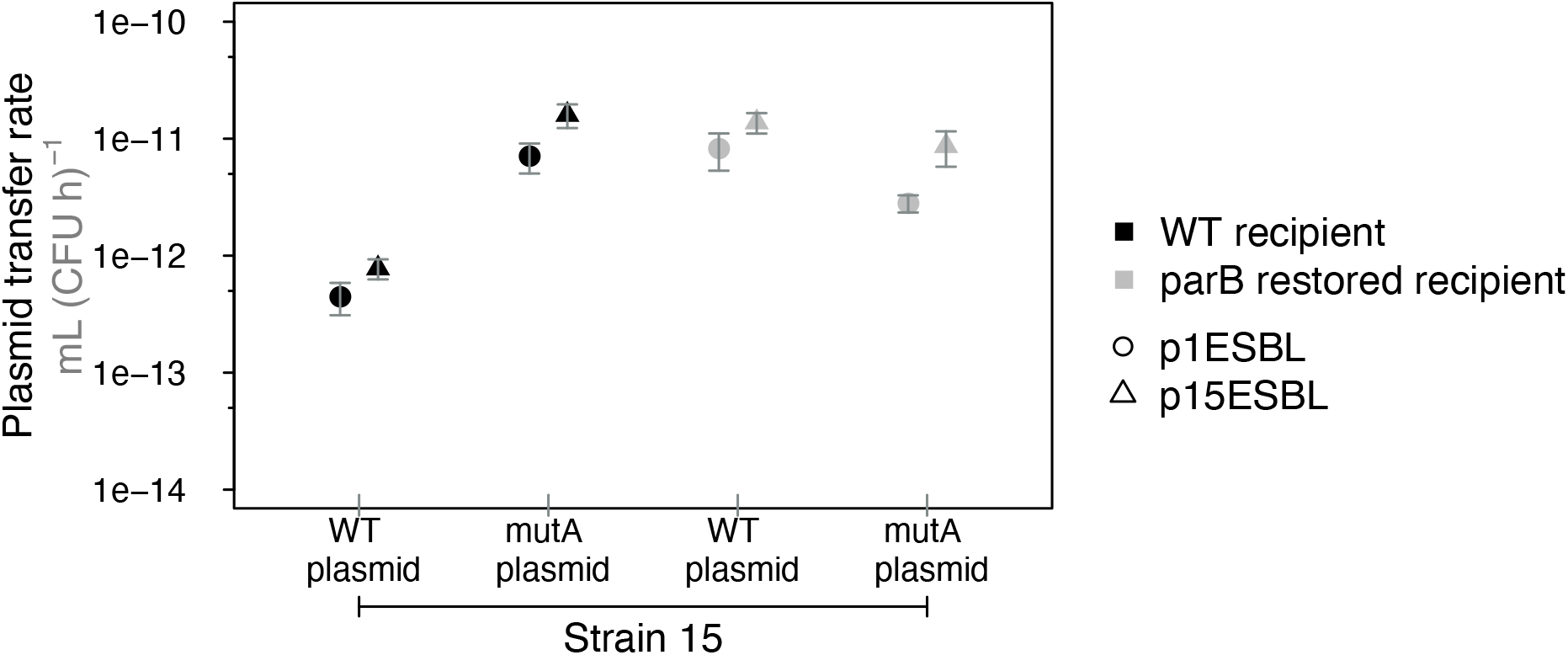
Complementing the *parB*-like protein in recipient strains did not restore ancestral plasmid transfer-rates. We conjugated wildtype (WT) and mutA (**Fig. 7B**) versions of plasmids p1ESBL and p15ESBL from strain 15 donors to marked strain 15 recipients, with or without complementation of the *parB*-like gene in the recipient strains. Complementation increased transfer rates for WT plasmids, but did not affect transfer of mutant plasmids (two-way ANOVA, complementation × strain (WT/mutA) interaction: *F* _1, 28_ = 18.434, *p* < 0.001; main effects NS). This suggests a positive effect of the *parB*-like protein on transfer, but only if the WT *ssi3* operon is present. The fact that this operon is largely conserved among conjugative plasmids (Wein *et al*., 2021) supports the idea of functional interaction among these genes.

### 4) Supplementary Tables

**Table S1:** Verification of ESBL-plasmids presence in three independent Amp resistant clones from each endpoint population. Primers used to verify plasmid presence/ absence. Yes/ no indicates successful/ unsuccessful amplification of indicated regions.

| Strain-plasmid pair | Population ID | Population colour | oriT amplicon | plasmid specific amplicon |
| --- | --- | --- | --- | --- |
| s1_p1 | 22 | green | yes | not tested |
| s1_p1 | 50 | orange | yes | not tested |
| s1_p1 | 53 | purple | yes | not tested |
| s1_p1 | 55 | pink | yes | not tested |
| s1_p15 | 2 | orange | yes | not tested |
| s1_p15 | 7 | pink | yes | not tested |
| s1_p15 | 29 | green | yes | not tested |
| s1_p15 | 40 | purple | yes | not tested |
| s15_p1 | 6 | orange | not tested | yes |
| s15_p1 | 39 | green | yes | yes |
| s15_p1 | 51 | purple | not tested | yes |
| s15_p1 | 43 | pink | not tested | yes |
| s15_p15 | 24 | orange | yes | no |
| s15_p15 | 27 | pink | yes | no |
| s15_p15 | 37 | green | yes | no |
| s15_p15 | 46 | purple | no | no |
| s19_p1 | 4 | purple | not tested | yes |
| s19_p1 | 17 | pink | not tested | yes |
| s19_p1 | 31 | green | not tested | yes |
| s19_p1 | 41 | orange | yes | no |
| s19_p15 | 1 | orange | not tested | yes |
| s19_p15 | 13 | green | not tested | yes |
| s19_p15 | 18 | purple | not tested | yes |
| s19_p15 | 26 | pink | not tested | yes |

**Table S2:**
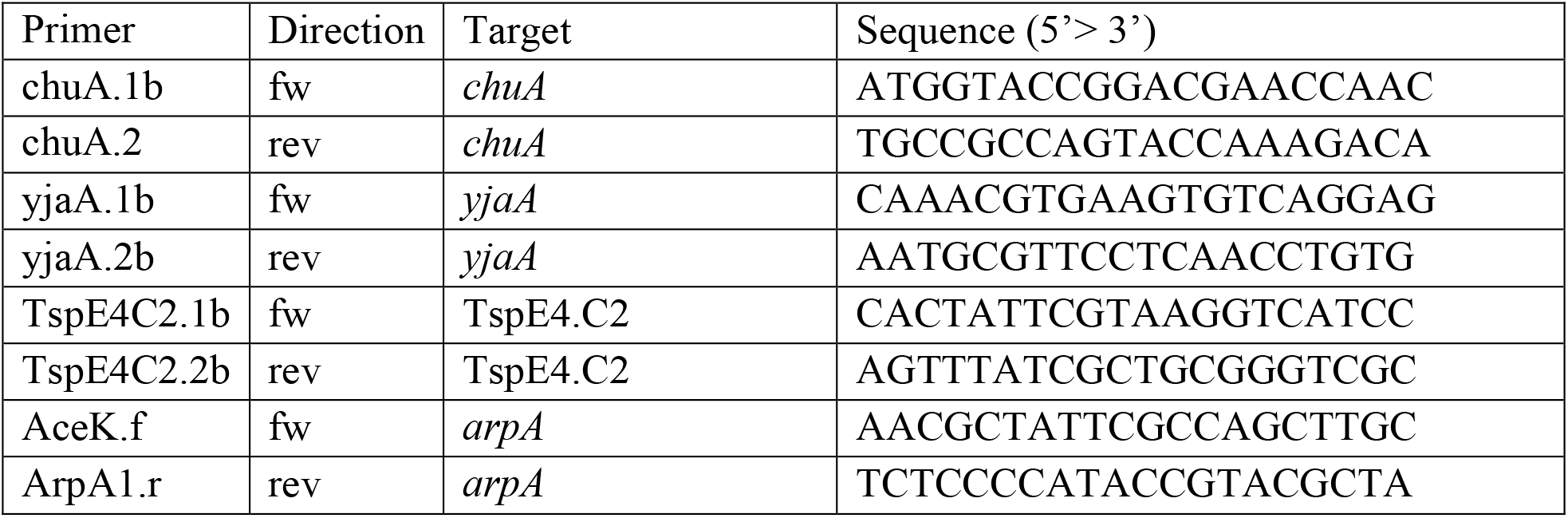
Primers used for quadruplex PCRs for Clermont phylo-typing to verify strains 1, 15, and 19 (see Supplementary Methods).

**Table S3:** Clinical *E. coli* strains (sequence type, ST, in brackets) and plasmids used to construct the strain-plasmid combinations used in this study. For each strain, each row gives one plasmid, identified by its replicon type(s). ESBL plasmids are indicated by strain-specific names in brackets; p1ESBL and p15ESBL are conjugative, p19ESBL is not and was cured from strain 19 to allow selection of p1ESBL and p15ESBL. To facilitate the introduction of p1ESBL and p15ESB into strain 19, IncIγ was also cured from strain 19. Resistance determinants: p1ESBL encodes for *aadA5, bla*_CTX-M-1_, *dfrA17* and *sul2*, p15ESBL for *bla*_CTX-M-1_ and p19ESBL for *bla*_CTX-M-27_, *tet(A), aph(6)-Id, aph(3’’)-Ib, sul2, mph(A), sul1, aadA5*, d*frA17, ant(3’’)-Ia*. With these three strains and p1ESBL and p15ESBL we created a fully factorial set of six strain-plasmid combinations. The star (*) with the IncFIC(FII) replicon in strain 1 indicates predictive uncertainty because the blast hit to IncFIC(FII) was shorter than < 75% coverage (Benz *et al*., 2020).

| Strain 1 (ST 117) |  | Strain 15 (ST 40) |  | Strain 19 (ST 131) |  |
| --- | --- | --- | --- | --- | --- |
| Replicon | Size (kb) | Replicon | Size (kb) | Replicon | Size (kb) |
| IncIγ (p1ESBL) | 111 | IncIγ (p15ESBL) | 88.9 | IncIγ (cured) | 87.7 |
| IncFIA;IncFIB; IncFIC(FII) | 153.9 | Col(MGD2); ColRNAI | 5.3 | IncFII;IncFIB; IncFIA;Col156 (p19ESBL) | 130 |
| IncFIC(FII)* | 86 | Col156 | 4.7 | Col156 | 5.2 |
| ColRNAI | 7 |  |  | Col(MG828) | 1.5 |

**Table S4:** Accessory genes of plasmid p1ESBL, generated with VrProfile (Li *et al*., 2018). The following features were tested: **VF**; Virulence factor, **AR**; Antibiotic resistance determinant, **T3SE**; (TE) type III secretion effector, **T4SE**; (FE) type IV secretion effector, **T6SE**; (SE) type VI secretion effector, **T7SE**; type VII secretion effector, **Prophage**; Prophage, **ICE**; integrative and conjugative element, **T3SS**; (T) type III secretion system, **T4SS**; (F) type IV secretion system, **T6SS**; (S) type VI secretion system, **T7SS**; type VII secretion system, **Integron**; Integron, **IS**; IS element, **PAI**; Pathogenicity island, **ARI**; Resistance island.

**Table S4** is provided in a separate file, SupplementaryDataFiles.xlsx, because it is too large to include here.

**Table S5:** Accessory genes of plasmid p15ESBL generated with VrProfile (Li *et al*., 2018). The following features were tested: **VF**; Virulence factor, **AR**; Antibiotic resistance determinant, **T3SE**; (TE) type III secretion effector, **T4SE**; (FE) type IV secretion effector, **T6SE**; (SE) type VI secretion effector, **T7SE**; type VII secretion effector, **Prophage**; Prophage, **ICE**; integrative and conjugative element, **T3SS**; (T) type III secretion system, **T4SS**; (F) type IV secretion system, **T6SS**; (S) type VI secretion system, **T7SS**; type VII secretion system, **Integron**; Integron, **IS**; IS element, **PAI**; Pathogenicity island, **ARI**; Resistance island.

**Table S5** is provided in a separate file, SupplementaryDataFiles.xlsx, because it is too large to include here.

**Table S6:** Description of genes in the leading region of p1ESBL and p15ESBL and their comparison to genes in the leading region of R64 (IncI1, accession AP005147) and pCT (IncK, accession FN868832). For genes indicated with a *, we performed functional predictions with Phyre^2^ homology modeling (Kelley *et al*., 2015).

**Table S6** is provided in a separate file, SupplementaryDataFiles.xlsx, because it is too large to include here.

**Table S7:** Accessory genes within the region of the putative chromosomal PAI of strain 19 (contig 1; position 1’777’892-1’932’181) generated with VrProfile (Li *et al*., 2018). The following features were tested: **VF**; Virulence factor, **AR**; Antibiotic resistance determinant, **T3SE**; (TE) type III secretion effector, **T4SE**; (FE) type IV secretion effector, **T6SE**; (SE) type VI secretion effector, **T7SE**; type VII secretion effector, **Prophage**; Prophage, **ICE**; integrative and conjugative element, **T3SS**; (T) type III secretion system, **T4SS**; (F) type IV secretion system, **T6SS**; (S) type VI secretion system, **T7SS**; type VII secretion system, **Integron**; Integron, **IS**; IS element, **PAI**; Pathogenicity island, **ARI**; Resistance island.

**Table S7** is provided in a separate file, SupplementaryDataFiles.xlsx, because it is too large to include here.

**Table S8:** Accessory genes within the region of the putative chromosomal PAI of strain 1 (contig ESBL01_6; position 19’420-40’408) generated with VrProfile (Li *et al*., 2018). The following features were tested: **VF**; Virulence factor, **AR**; Antibiotic resistance determinant, **T3SE**; (TE) type III secretion effector, **T4SE**; (FE) type IV secretion effector, **T6SE**; (SE) type VI secretion effector, **T7SE**; type VII secretion effector, **Prophage**; Prophage, **ICE**; integrative and conjugative element, **T3SS**; (T) type III secretion system, **T4SS**; (F) type IV secretion system, **T6SS**; (S) type VI secretion system, **T7SS**; type VII secretion system, **Integron**; Integron, **IS**; IS element, **PAI**; Pathogenicity island, **ARI**; Resistance island.

**Table S8** is provided in a separate file, SupplementaryDataFiles.xlsx, because it is too large to include here.

**Table S9:** Comprehensive list of genetic changes identified comparing ancestral with evolved clones. Non-self-explanatory column names: Pop ID = Identifier of the population the clones were isolated from; Clone ID = Clone specific identifier (p+/p-); Col = Colour, Contig = Contig according to deposited sequences; Del start/ end = Start and stop of deletions larger than 10 bp.

**Table S9** is provided in a separate file, SupplementaryDataFiles.xlsx, because it is too large to include here.

**Table S10:**
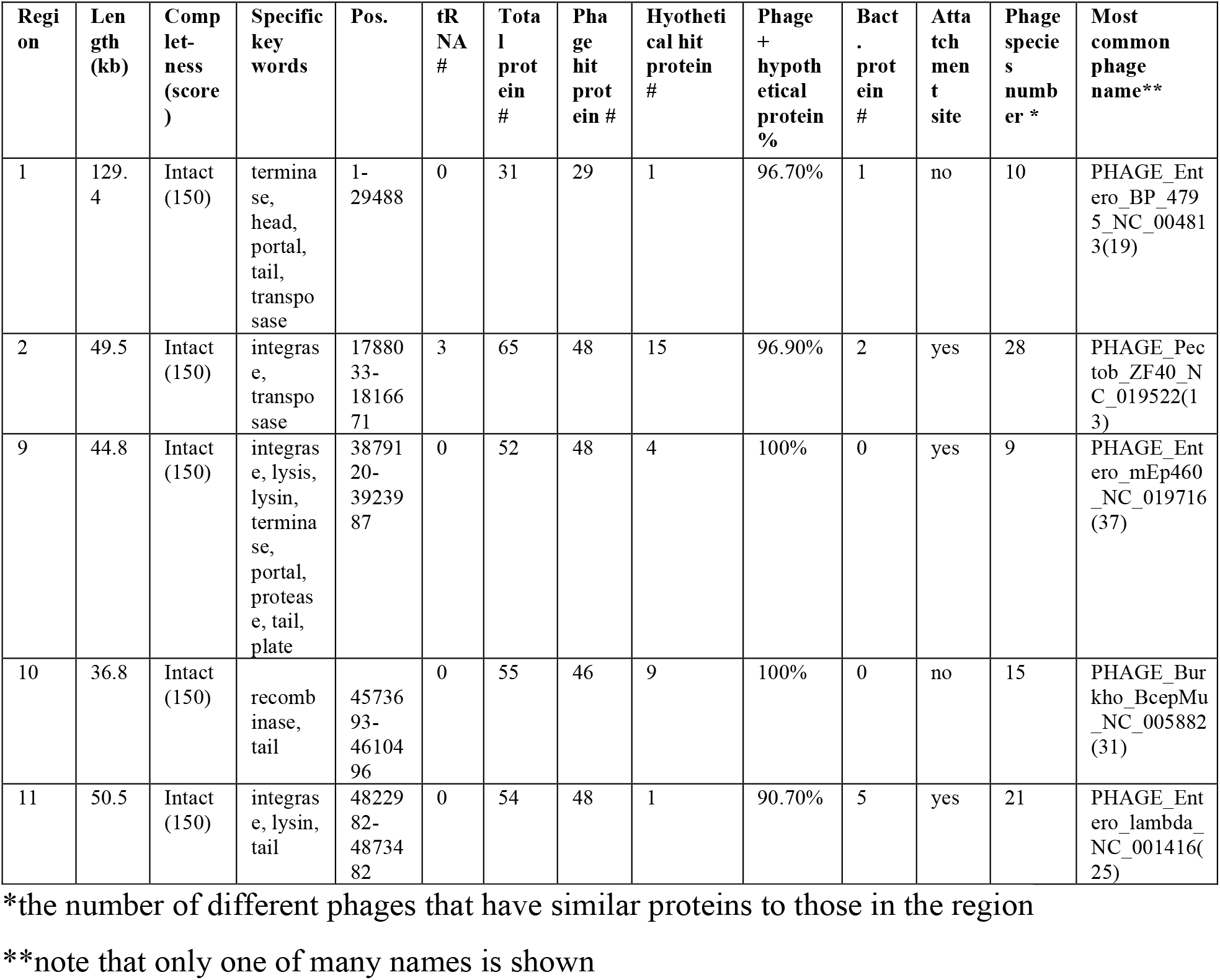
Phage prediction for strain 19, performed with Phaster (Arndt *et al*., 2016). Note that only the first of the most common phage names are given. Phages present in strain 1 and 15 are presented in Benz et al. (2020).

**Table S11:**
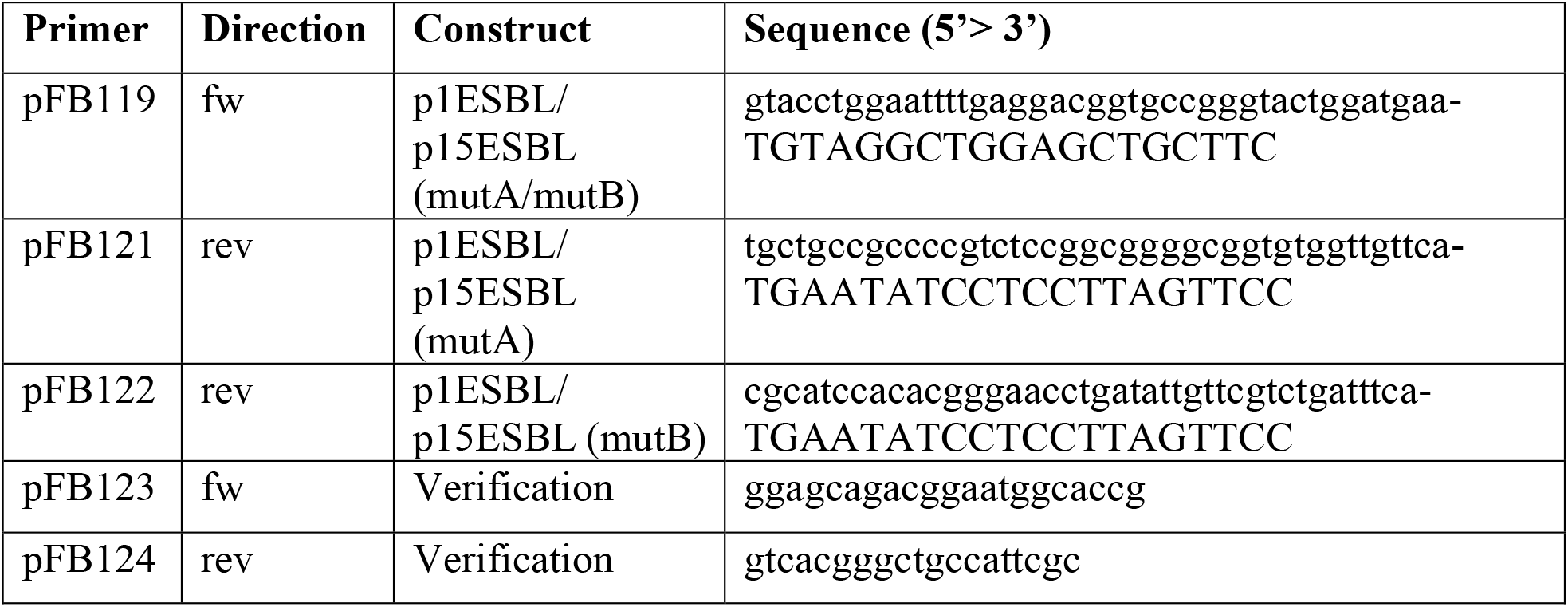
Primers used to generate knockout mutants of p1ESBL and p15ESBL.

